# Advancing genetic engineering in the plant growth-promoting agricultural bacterium *Azospirillum brasilense*

**DOI:** 10.1101/2025.04.08.647883

**Authors:** Emma Riley, Melissa Poma, Ciarán L. Kelly

## Abstract

*Azospirillum brasilense* is an important plant-growth promoting bacterium found in the rhizosphere and used extensively in commercial agriculture. It is an attractive candidate for use in the development of novel crop interventions, such as the targeted delivery of plant hormones and similar compounds to the roots, shoots or leaves, or the introduction of smart bacterial sensors and actuators in the rhizosphere. In order to be able to engineer *A. brasilense* with these novel and complex behaviours, we require a collection of predictable and reliable genetic parts, consisting of constitutive and inducible promoters, terminators, and a stable expression plasmid. To date, such a genetic toolkit does not exist for the genus. In this work, for the first time we have designed and tested a synthetic constitutive promoter library with a wide-range of transcriptional strengths, a tightly-regulated inducible promoter with a non-metabolisable inducer, strong Rho-independent transcriptional terminators and synthetic small RNAs for post-transcriptional regulation.. We have adapted them for use in a one-pot assembly method and demonstrated their utility in the overproduction of the important plant hormone, indole 3-acetic acid (IAA).

## Introduction

The resilience of crop production is an issue of urgent worldwide concern. It is predicted that crop production must increase by 70% by 2050 to meet the growing food demand of the global population ^1,2^. At the same time, adverse environmental conditions such as extreme heat, flooding, and drought, threaten the stability of our ecosystems and agricultural productivity ^3^. Climate change has already resulted in more frequent and severe plant pathogen outbreaks in areas with elevated temperatures, higher carbon dioxide levels and soil moisture content ^4^.

Predicted increases in global temperatures of 2 - 4 °C over the next century could result in a 6 - 18% reduction in global crop yields ^2,5^. Modern agricultural practices to boost crop yields, include the application of high concentrations of artificial nitrogen and phosphorus fertilisers to the soil, with the concurrent spraying of pesticides. Unfortunately, the increase and reliance on artificial fertilisers and pesticides is directly responsible for extensive ecological damage, including eutrophication and hypoxia of waterways and biodiversity loss^6–9^, as well as damage to human health from particulate air pollution through the combination of farmland ammonia with traffic emissions^10^. Additionally, widespread pesticide use has led to the evolution of resistance amongst pathogens, with many fungal pathogens resistant to all commonly-used fungicides ^11^, compounded further by recent evidence that nitrogen and phosphorus fertilisers encourage the growth of pathogenic fungi over mutualistic fungi ^12^.

Agricultural biologicals are natural products and microbial bioinoculants that are added to plant seeds and soil as (a) biofertilisers fixing atmospheric nitrogen, (b) biocontrol agents against pathogens, or (c) biostimulants for enhanced plant growth. They had a global market value of $13.76 billion in 2023 and this is expected to triple by 2035 to $43.53 billion. Microbial bioinoculants typically consist of formulations of one or more species of plant-growth promoting (PGP) bacteria and are growing in popularity as an environmentally-friendly intervention to boost crop productivity ^13^. Many plant growth-promoting rhizobacteria with a variety of plant association mechanisms have been described. Some remain free-living in the soil and colonise the surface of roots. Others, such as rhizobia enter the superficial spaces of plants such as legumes and form root nodules. Others still can grow in the space between the plant’s cell wall and plasma membrane as obligate or facultative endophytic bacteria ^14^. Motivated by the need to reduce our reliance on artificial fertilisers, the engineering of biological nitrogen fixation is a huge area of current synthetic biology research ^15–20^. Others strive to achieve ‘’artificial symbioses’ in non-leguminous cereals but research is limited by the genetic and regulatory constraints of transgenic plants ^7,15^.

Members of the genus *Azospirillum* are some of the best characterised plant-growth-promoting rhizobacteria. *Azospirillum* species are found in almost all soil types on Earth and can colonise over 100 plant species, including economically-important crops such as maize, wheat, and barley. They have been shown to enhance plant growth by 10-15% through multiple mechanisms working in combination ^1,14,21^. These include nitrogen fixation and the solubilisation of inorganic phosphates (biofertiliser activity); the production of a variety of chemical compounds such as auxins and hormones to stimulate changes in root growth (biostimulant activity) ^22^; and protection from both bacterial and fungal pathogens (biocontrol activity) ^23,24^. As *Azospirillum* is one of only four bacteria approved as biofertilisers by the European Union, it is a strong candidate for engineering, for example to enhance nitrogen fixation in different environmental conditions, to increase production of specific PGP chemicals for use with specific plants, to boost its biocontrol activity, or to introduce new complementary plant-growth promoting capabilities to the rhizosphere. Despite some basic genetic studies looking at colonisation mechanisms, and the regulation of the important indole 3-acetic acid (IAA) pathway ^25–28^, few genetic tools have been developed for any *Azospirillum* species. Previous attempts to engineer *Azospirillum* strains relied upon a handful of native constitutive promoters and promoters previously used in common *Eschericia coli* (*E. coli*) vectors ^29,30^.

In this work, for the first time, we report a suite of genetic tools for the high-resolution control of transcription and translation in *Azospirillum brasilense* sp7, a well-studied representative member of this important genus that has been used in commercial farming for many years ^21^. This toolkit introduces new control and tunability of gene expression and protein production for the engineering of *Azospirillum brasilense* sp7, both for fundamental studies as well as novel agritech applications. We demonstrated the power of the new genetic toolkit for the successful generation of indole 3-acetic acid (IAA) overproduction strains.

## Results + Discussion

### A stable origin of replication plasmid for the engineering of *A. brasilense*

Various methods for chromosomal integration in *Azospirillum* have been described, but to facilitate high-throughput engineering, we wanted to identify a multicopy plasmid with stable replication in lab conditions. Two plasmids with commonly-used origins of replication were tested, one containing the p15A origin, commonly used in *E. coli,* and the other containing the pBHR1 origin, a broad-host range origin, and a gene encoding the mobilisation protein, *mob* ^31,32^. In brief, *A. brasilense* sp7 *(A. brasilense* hereafter) cells were transformed by electroporation with either the p15A or pBHR1 plasmid and transformants sequentially subcultured on agar plates with or without appropriate antibiotics. Plasmid stability was determined by the ability of colonies from agar plates lacking antibiotic to subsequently grow on antibiotic-containing agar plates, thus indicating the maintenance of the plasmid conferring antibiotic-resistance cassette. The majority of cells (96.6%) transformed with the p15a-derived plasmids had lost resistance after the first passage (25 generations) and had lost all resistance after the second passage (Figure S1). In contrast, we found the pBHR1-derived plasmid to be stably maintained in the absence of that selective pressure, with no statistically-significant loss of resistant cells over the course of the five successive passages (Figure S1). This pBHR1-derived plasmid was used for all subsequent experiments in both *A. brasilense* and *E. coli*.

### A set of species-specific constitutive promoters with a wide range of transcription strength

As a limited number of promoters, representing a narrow range of expression levels, had previously been used in *Azospirillum*, a synthetic promoter library (SPL) was generated. To ensure promoters with a wide range of expression were constructed, which would allow control and tuning of transcription strengths, and avoid homologous recombination of promoter libraries derived from single-base pair mutation libraries of an existing promoter, the Jensen and Hammer ^33^ method was chosen. All putative sigma 70 promoters in the sp7 genome were identified using the BPROM prediction tool ^34^, and any sequences located in open reading frames were omitted using a custom Python script. The-35 and-10 nucleotide sequences were identified and consensus sequences for each region determined (Figure 1A). With the relative nucleotide frequencies for the-35 and-10 promoter sequences in *A. brasilense* sp7 obtained, we determined a minimum threshold for the frequency of each nucleotide in each sigma factor binding regions of 0.6, with nucleotides over this value conserved in the library design. Where a single base did not meet this minimum threshold level, the second or third most common nucleotide was included until the threshold was reached. The final conserved sequences were TTGMMD (-35) and YANNMT (-10), where M represents A or C; D represents A, G, or T; Y represents C or T; and N represents A, C, T, or G (Figure 1B). Finally, the 33 nucleotides surrounding these conserved regions were then randomised giving a theoretical library size of approximately 7.4 x 10^19^. These promoter sequences were introduced upstream of a gene encoding GFP on the pBHR1 plasmid backbone (Figure 1C). *A. brasilense* sp7 cells were transformed with this library and 94 single colonies chosen at random, cultured in microplates until mid-exponential phase of growth, and GFP fluorescence measured using a microplate reader to determine the range of expression strengths present in the library (Figure S2). To examine promoter sequence diversity and structure, forty clones were taken forward to test in triplicate in fluorescence assays by flow cytometry.

**Figure 1.**
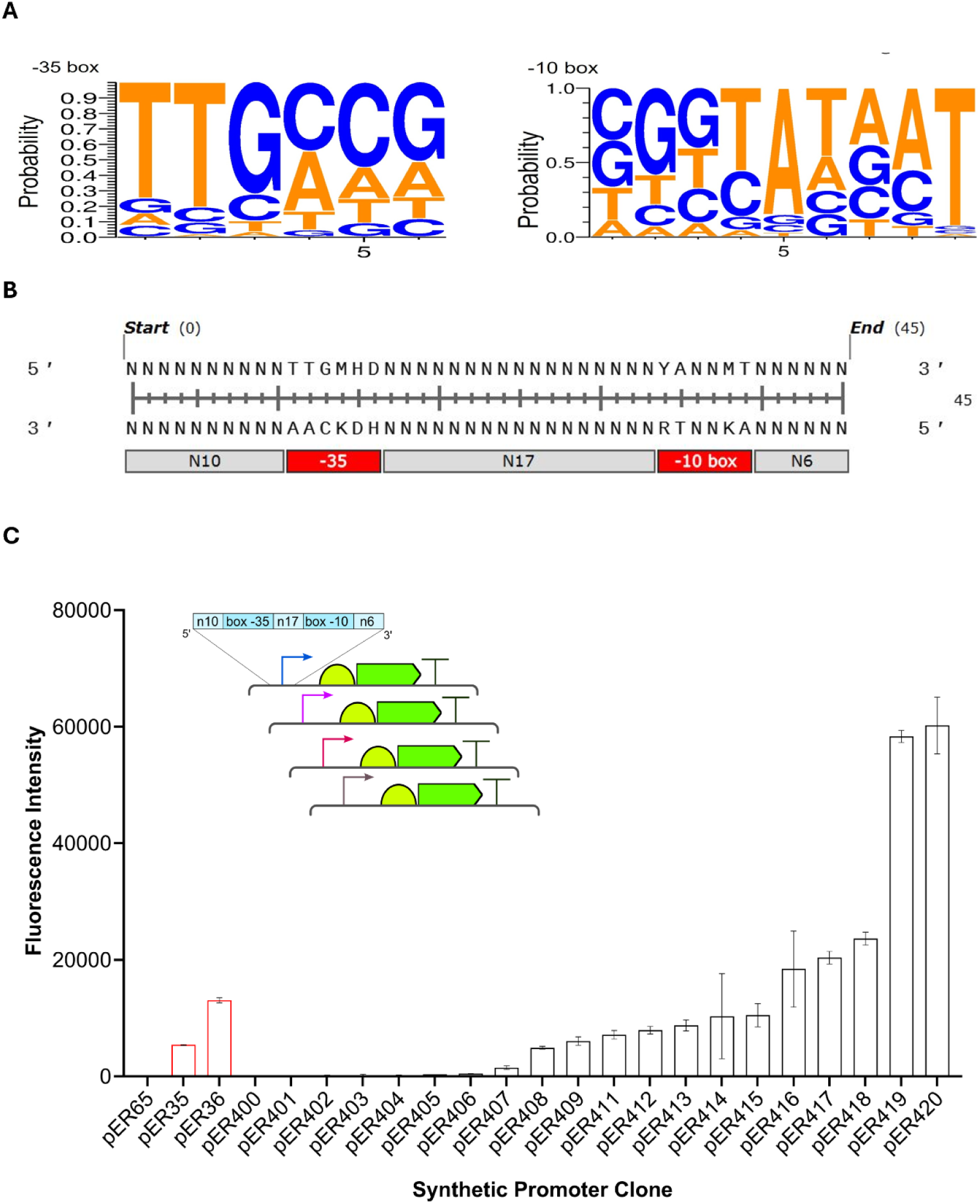
A synthetic promoter library (SPL) for A. brasi/ense sp7. A) Identification of the consensus sigma70 promoter sequences for A *brasilense* sp7. The A *brasilense* sp7 reference genome NZ_CP012914.1 was analysed using BPROM to predict promoter sequences. A custom Python script was used to decode the-35 and-10 promoter boxes from the ORF-less genome and-35 and-10 consensus promoter sequences visualised using Weblogo^37^. B) Final synthetic promoter sequences used for SPL generation. Mutagenic forward and reverse primers were designed to complement the consensus synthetic promoter sequence, incorporating nucleotide ratios of N = 25% G, A, T, C; D = 33.33% A, G, T; M = 50% **A,** C; Y = 50% C, T and conserving bases identified by the consensus promoter analysis C) Design of SPL screen (inset) and flow cytometry data for SPL. Characterisation of twenty synthetic promoter clones isolated from the initial library screen. A *brasi/ense* sp7 cells containing SPL plasmids were isolated and fluorescence assayed by flow cytometry. Plasmids pER35 and pER36 represent the reference BBa_J23118 and ProD promoter controls respectively and pER65 is a no promoter negative control. Fluorescence intensity was measured following 8 hours of growth in MMAB. Error bars shown represent the standard deviation of the mean of three independent biological replicas.

A plasmid lacking a promoter, pER65, and two reference promoter plasmids, pER35 (containing the medium-strength Anderson promoter BBa_J23118 ^35^) and pER36 (containing the strong synthetic promoter proD ^36^ were included as controls. Transformants displaying multimodal fluorescence were discarded, with twenty plasmids remaining. A wide range of expression strengths were observed in the library, with weak, medium, strong and very strong promoter activities observed relative to the reference promoters (Figure 1C).

As pER419 and pER420 were almost three times greater than the nearest promoter in strength, we tested the residuals for normal distribution. A one-way ANOVA model was fitted, the residuals extracted, a histogram and Q-Q plot generated (Figure S3) and the residuals analysed using the Shapiro-Wilk test for normal distribution. As expected the histogram showed a skewed distribution with some extreme values, especially on the right; the Q-Q plot showed residuals deviated noticeably from the straight line, especially in the tails, suggesting non-normality, and the Shapiro-Wilk test resulted in a p-value of 2.46 × 10⁻¹¹ confirming the residuals are not normally distributed. To overcome this we transformed the data into log_10_ values to obtain a normal distribution (Figure S4), and a one-way ANOVA statistical test was conducted, showing a statistically-significant difference in the means across the promoter set (P = <0.0001). Individual t-tests with Tukey’s corrections for multiple comparisons were conducted to assess pairwise differences between promoter means. All synthetic promoters except pER400 showed significantly-different fluorescence to the negative control plasmid pER65. The results were summarized using a compact letter display, where promoters sharing a letter are not significantly different (α = 0.05), while groups without shared letters are distinct (Figure S5). Several promoters exhibited overlapping group labels (e.g., pER417 labeled “C D”), indicating that their expression was statistically indistinguishable from both C and D groups. This overlap suggests subtle gradations in expression strength, which allows for high-resolution promoter selection from 11 statistically-distinct expression tiers (A–K), demonstrating that the synthetic promoter library achieves fine-grained control over gene expression levels — far beyond simple low/medium/high categorisation. This supports the library’s utility for precise tuning of biological circuits and expression systems.

Following the analysis of expression strengths by flow cytometry, the promoter sequences were determined by Sanger sequencing and aligned using a forced motif-anchored method. The closest match to the TTGMMD consensus for the-35 box was identified in each sequence using degenerate IUPAC matching with tolerance to mismatches. The-10 box (YANNMT) was estimated 17 bases downstream of the-35 box when it could not be confidently matched. This approach allowed all promoter variants to be included in the alignment while preserving the functional layout of core elements (-35 and-10 boxes) (Figure S6). Some loss of conserved motifs within the-35 and-10 elements were observed, which could result from errors during oligonucleotide synthesis or machine mixing or artefacts of blunt-end ligation cloning method. A custom Hamming distance was calculated between all aligned promoter sequences, excluding positions with alignment gaps. The distance reflects the proportion of mismatched nucleotide positions among valid comparisons. The distance matrix was standardised and subjected to PCA using the scikit-learn library. Expression bins were assigned from an updated classification set and used to annotate promoter clusters in PCA space. Promoters were grouped into refined expression bins including’low’,’mid-low’,’medium’,’mid-high’,’strong’, and’very strong’ (Table S6). The PCA revealed a broad and structured distribution of promoter variants across principal components (Figure S7), reflecting meaningful sequence diversity. The clustering showed that high-and low-expression promoters were not isolated in a single cluster, supporting the presence of sequence-driven functional variation. The overlap and gradients observed suggest nuanced regulatory influence encoded in the promoter variants.The first two principal components were plotted to visualize major dimensions of sequence variation across the promoter set.

### An inducible promoter system for temporal control of transcription

To complement this suite of constitutive promoters, we wanted to identify an inducible promoter system for use in *A. brasilense* sp7. Unlike constitutive promoters with fixed strengths of transcription, inducible promoters allow the strength and timing of gene expression to be controlled using one DNA sequence and the addition of an exogenous signal (inducer). None have been described previously for use in *A. brasilense* sp7. The properties of an ideal inducible promoter would include: no basal expression (“leakiness”) in the absence of inducer, which is important when gene product creates cellular burden and/or toxicity; a linear response of expression to increasing concentrations of inducer, allowing precise control of expression levels; and use a non-toxic, non-metabolised inducer, ensuring sustained expression during culturing. Three commonly-used inducible promoter systems were assessed for use: the allolactose/IPTG-inducible promoter P*_lac_*; the tetracycline/anhydrotetracycline-inducible promoter P*_tet_*; and the L-rhamnose-inducible promoter P*_rhaBAD_*. The promoter sequences were inserted in place of the constitutive promoters on the previously-used GFP-reporter plasmid pER35. Homologues of the genes encoding the promoter-associated transcription factors, LacI and TetR were identified on the *A. brasilense* sp7 genome through BLAST searches and thus not added to reporter plasmids. No gene encoding the homologue of the activating transcription factor of P*_rhaBAD_*, RhaS was identified, so an additional plasmid was constructed where the *rhaS* gene and its native RBS was inserted downstream of the *kanR* gene on pBBR1, as previously done (Figure 2A)^37,38^. No induction was observed in *A. brasilense* sp7 cells containing either the P*_lac_* or P*_tet_* promoter-GFP plasmids with any concentration of IPTG or anhydrotetracycline. Cells transformed with the P*_rhaBAD_* promoter-GFP plasmid showed a linear response of fluorescence to the concentration of L-rhamnose added, when sampled after 8 h of culturing (late exponential phase of growth), for both pER70 (+*rhaS*) and pER72 (-*rhaS*) (Figure 2B). To determine if the slopes of the linear regressions differed significantly between pER70 and pER72, a linear regression was performed (excluding the 0 mg/ml L-rhamnose data points) and a two-tailed t-test was performed. A p-value of 0.139 indicated no statistically significant difference in slopes and thus no difference in the response to L-rhamnose of plasmids with or without *rhaS*. It was surprising that despite the lack of identification of a gene encoding a RhaS homologue, the absence or addition of the constitutively-expressed *rhaS* had no effect on induction. Finally, growth assays were set up in minimal media supplemented with either malate or L-rhamnose to assess if L-rhamnose was metabolised by *A. brasilense* sp7, which would lead to a reduction in induction of any target gene over time. No growth was observed with L-rhamnose as the sole carbon source (Figure S8).

**Figure 2.**
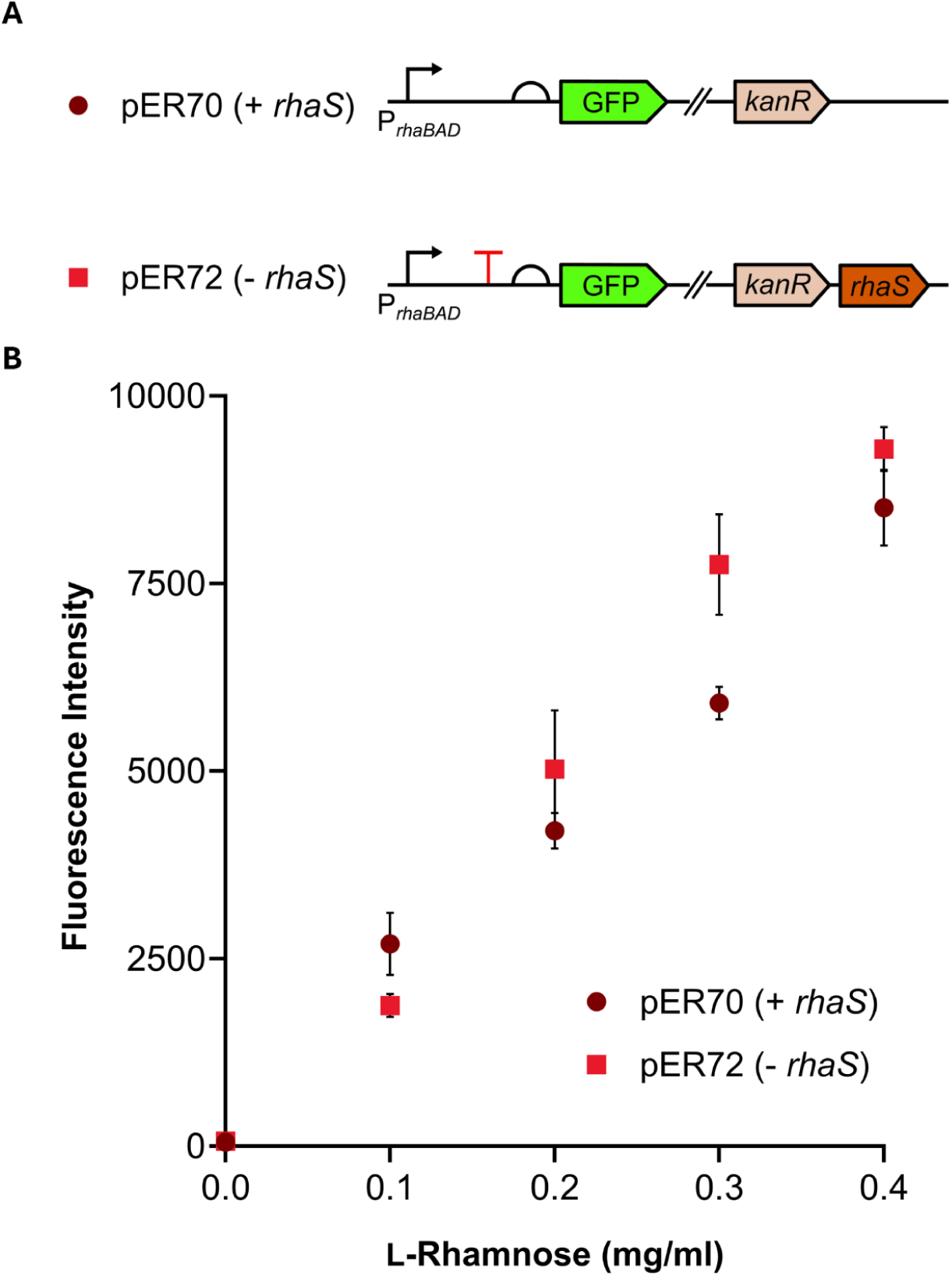
Testing of an L-rhamnose inducible promoter system in A. brasilense sp7. A) Design of reporter plasmids pER7O (with heterologous constitutive expression of *rhaS* from *E. col!)* and pER72 (no *rhaS).* B) Testing of promoter response to increasing concentrations of L-rhamnose. *A. brasilense* sp7 transforrnants containing plasmids pER70 or pER72 were cultured in MMAB media supplemented with increasing concentrations of L-rha mnose. Fluorescence intensity was measured by flow cytometry following 8 hours of growth. Error bars shown represent the standard deviation of the mean of three independent biological replicas.

### Transcriptional terminators to achieve independent control of gene expression for multi-enzyme pathway engineering

Rho-independent or intrinsic transcriptional terminators involve the formation of a hairpin loop in the RNA secondary structure as the RNA polymerase extends the nascent mRNA strand. This hairpin loop causes dissociation of the RNA polymerase and thus halts RNA elongation and thus transcription. Many intrinsic terminators have been characterised in other organisms and they are essential for the construction of complex multi-gene systems,allowing the isolation or insulation of one transcriptional unit from the next, and different promoters with different transcriptional strengths to be tested to find the optimal expression level of each component in the system. No terminators have ever been tested in *Azospirillum* to date, preventing the ability to quickly build and test new metabolic pathways or genetic circuits in this organism. We chose seven terminators: three native *E. coli* terminators and four synthetic terminators that had been shown to perform well in a variety of organisms with Gibbs free energies below-10 kcal/mol. Terminators were placed between the L-rhamnose inducible promoter and the ribosome binding site of the GFP reporter gene on pER70 (Figure 3). *A. brasilense* cells were transformed with each plasmid alongside no promoter (pER65) and no terminator (pER70) controls and cultured with or without L-rhamnose and GFP production tested by flow cytometry. No statistically-significant difference in fluorescence was observed between samples lacking inducer and those containing the inducer for all seven terminator-containing plasmids (Figure 3), indicating that all sequences tested provided robust transcriptional termination in *A. brasilense*.

**Figure 3.**
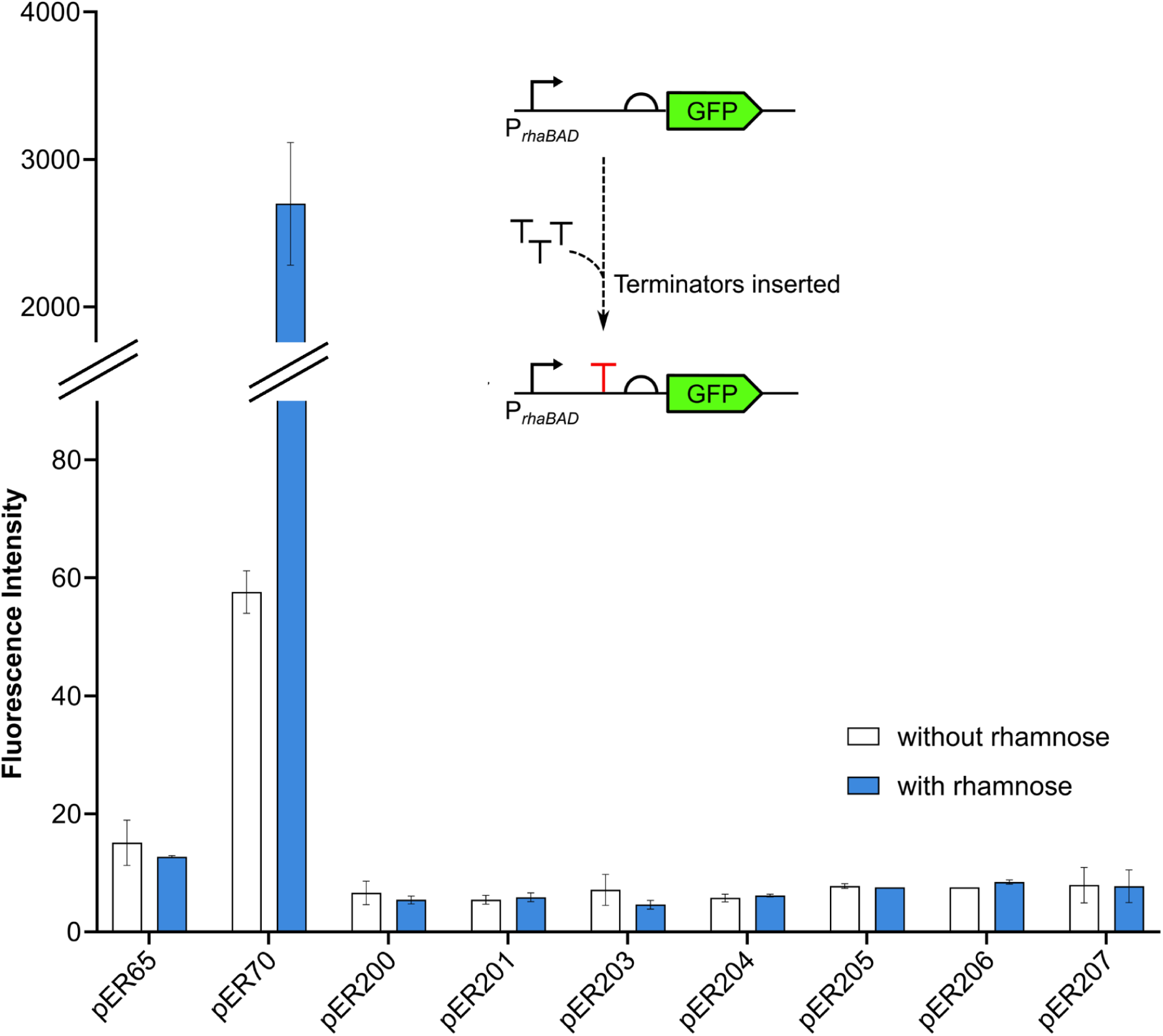
Screening of intrinsic transcriptional terminators in A. brasilense sp7. Design of the screen to test for effective termination of transcription (inset). Flow cytometry data for terminator screen with and without L-rhamnose. ***A.*** *brasilense* sp7 transformants of plasmids containing the appropriate terminator inserted between the L-rhamnose inducible PrhaBAD promoter and the ribosome binding site of the sfGFP reporter gene, no promoter (pER65), or no terminator (pER70) controls, were cultured in MMAB media supplemented with 0.4 mg/ml L-rhamnose. Fluorescence intensity was measured by flow cytometry after 8 hours. Error bars shown represent the standard deviation of the mean of three independent biological replicas.

### Synthetic sRNAs for post-transcriptional regulation in *A. brasilense*

Small bacterial RNAs (sRNAs) are short (approximately 200 bp), non-coding RNAs that anneal to target messenger RNAs (mRNAs) through complementary base pairing, aided by a protein chaperone Hfq. In the majority of cases reported, this causes translation inhibition of the target mRNA. They have many advantages over other gene-silencing mechanisms, being fast-acting and having a linear dose response due to the one-to-one stoichiometry of sRNA to mRNA. They are highly engineerable and specific, through the inclusion of a 24 bp seed region with reverse complementarity to the target sequence. Finally, as many Gram-negative bacteria encode a Hfq protein, they impart a much lower burden cost on host cells than approaches such as CRISPR that require heterologous protein production. Expanding the genetic toolkit for *Azospirillum* engineering to post-transcriptional regulation through synthetic sRNAs, introduces the ability to construct genetic circuits with greater layers of complexity for the first time, such as feedback circuits, logic gates and switches. Their successful introduction would also allow the rapid screening of gene-deletion candidates for metabolic engineering, for example to overproduce important plant hormones, without the time-consuming process of making knock-out strains.

Homology searches identified a homologue of the *E. coli* Hfq protein in the *A. brasilense* sp7 genome with 36.6% identity 51.5% similarity (Figure S8), suggesting synthetic sRNAs designed using previously-described principles^39^ should function in *A. brasilense*. To quickly screen for synthetic sRNAs that gave good translational inhibition, a dual sRNA-reporter plasmid system, pER100, was designed and constructed (Figure 4). The sRNA expression module comprises the L-rhamnose inducible promoter from pER70, a 24 bp seed region that can be changed to allow silencing of any mRNA sequence, the Hfq-binding scaffold from the *E. coli* micC sRNA, and a strong transcriptional terminator. The reporter module consists of a strong constitutive Anderson promoter (BBa_J23118) in front of the target reporter fragment, which is made up of the 30 bp upstream of the target open-reading frame, and the first 30 bp of the target orf itself, fused in frame with a gene encoding GFP lacking its start codon. This results in a translational fusion of the target gene product to GFP, allowing a simple fluorescent readout for successful inhibition of translation. SapI sites with unique overhangs were inserted around each module and *sacB-* and *lacZ-*expression cassettes introduced for rapid screening of successful sRNA and reporter cloning respectively. Cells containing a fully assembled sRNA-reporter screening pair can be screened for sRNA inhibition of translation, by testing for decreasing GFP fluorescence with increasing concentrations of L-rhamnose.

**Figure 4.**
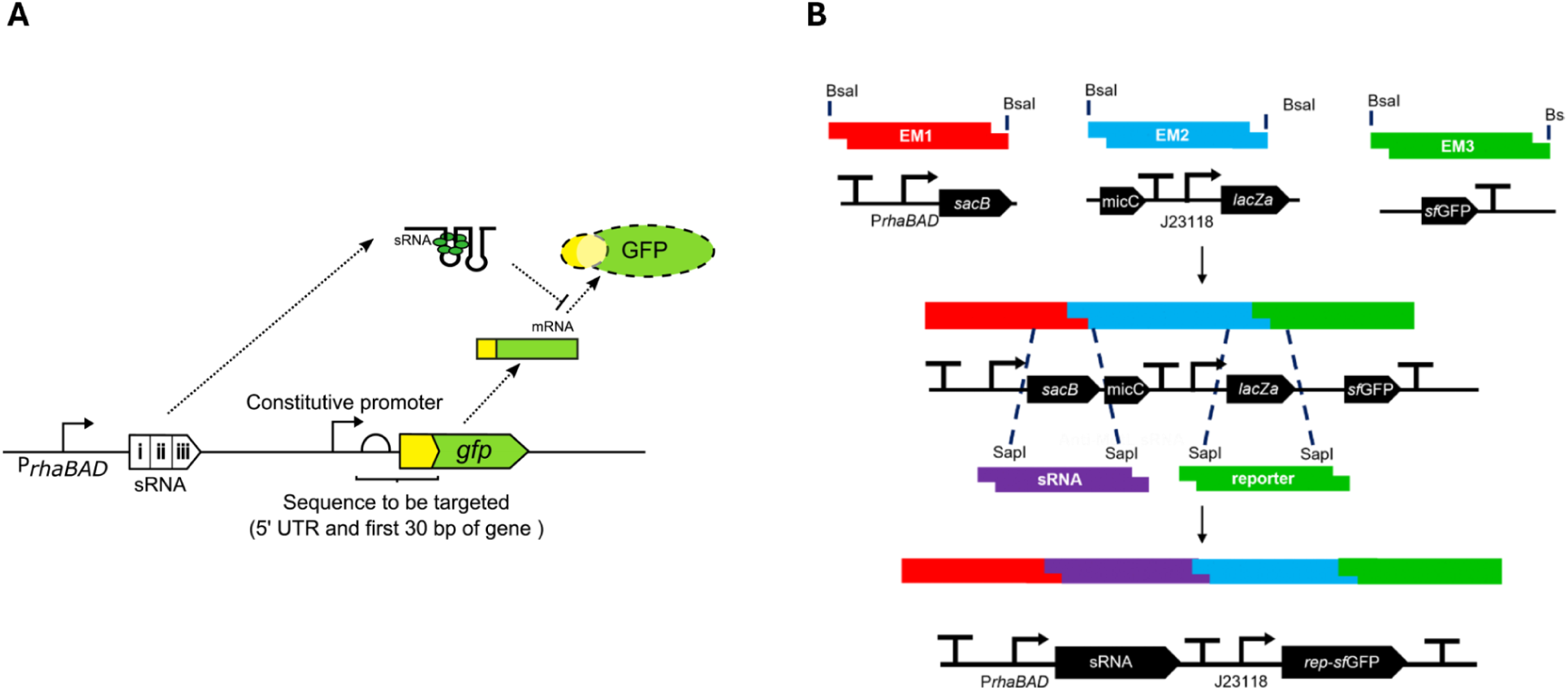
A dual sRNA and reporter screening plasmid to test for translation inhibition in A. brasilense. A) Design of sRNA and reporter screen. The inducible PrhaBAD promoter controls the expression of synthetic sRNAs that bind to constitutively-expressed target-reporter fusion mRNAs. Synthetic sRNAs are composed of a 24 bp mRNA target-binding sequence, a Hfq-binding scaffold sequence derived from the *E. coli* micC sRNA, and a strong transcriptional terminator. For screening and validation of sRNAs against mRNA targets, reporters were constructed containing the 30 bp located immediately upstream of the target gene and the first 30 bp of the target open­reading frame, which is fused to a gene encoding GFP. B) A one-pot assembly method for sRNA and reporter sequence cloning. The level 1 expression vector (pER100) was constructed with four unique Sapl restriction sites inserted either side of an sRNA-and reporter-sequence insertion site. To allow for easy identification of successful sRNA and reporter clones, a *sacB* and *lacZ* expression cassette was inserted in sRNA and reporter landing-spots respectively, to allow selection on sucrose agar plates and blue-white screening. sRNA and reporter sequences can then be cloned into level 0 holding vectors and combined with pER100 for one-pot assembly.

As an initial proof-of-principle, a native *A. brasilense* mRNA target was chosen, *mutL*, encoding a key protein in the methyl-directed mismatch repair (MMR) proteins. sRNA sequences were designed to target these and the 24 bp seed region analysed using IntaRNA^40^ to predict the binding energies of each sequence. Scores below-35.00 kcal/mol^-1^ were used as a threshold for strong predicted sRNA-mRNA binding. Highly favourable overall Gibb’s free energy scores were predicted for anti-*mutL* (-36.91 kcal/mol^-1^). This seed sequence and corresponding 60 bp reporter sequences were cloned via level 0 holding plasmids, into pER100, sequence verified, and subsequently introduced into *A. brasilense*, cultured in minimal MMAB media supplemented with varying concentrations of L-rhamnose, and the changes in fluorescence measured by flow cytometry at late exponential phase of growth. A reduction in fluorescence was observed with the anti-*mutL* plasmid, with increasing concentrations of L-rhamnose (Figure 5A), indicating successful translation inhibition was occuring in *A. brasilense*. However, even at appropriately high concentration of L-rhamnose, complete silencing of reporter-GFP production was not observed. We hypothesised that this was as a result of an imbalance between the numbers of sRNA and mRNA molecules present in the cell, leading to unbound mRNA molecules being free for translation initiation. It is likely that the combination of a constitutive promoter used to express the reporter-GFP fusion mRNA and the presence of the chromosomally-encoded mRNA target acting as a sponge for the sRNA molecules leads to a dampening of reporter-GFP mRNA inhibition.

**Figure 5.**
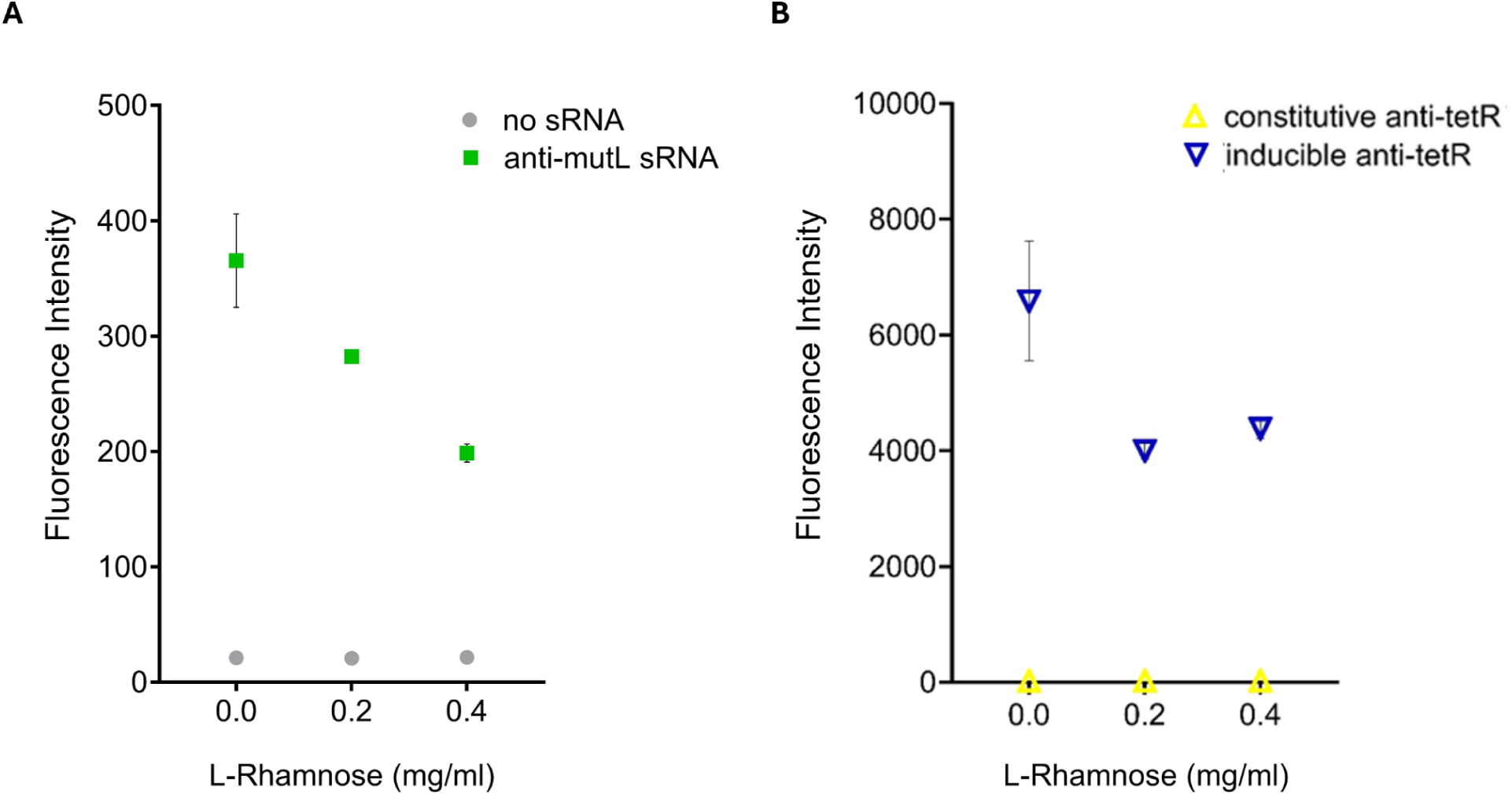
Testing synthetic sRNAs against mutL-and tetR-fused GFP reporters in A. brasilense. A) Testing of an anti-muL sRNA under the control of the L-rhamnose inducible promoter. *A. brasilense* sp7 cells were transformed with pER165 (anti-mutL/mutL-sfGFP) and single colonies grown at 30 °C overnight in MMAB media. Overnight cultures were then pre­cultured for 6 hours and normalised to a starting OD of 0.05 in fresh MMAB supplemented with 0.0, 0.2 and 0.4 mg/ml L-Rhamnose respectively, cultured for 8 hours and fluorescence measured by flow cytometry. B) Testing of inducible and constitutively-expressed anti-tetR sRNA. *A. brasilense* spľ cells were transformed with pER158 (inducible anti-tetR/tetR-sfGFP) and pER303 (constitutive anti-tetR/tetR-sfGFP) and single colonies were grown overnight in MMAB media at 30 °C. Overnight cultures were pre-cultured for 6 hours and normalised to a starting OD of 0.05 into fresh MMAB supplemented with 0.0, 0.2 and 0.4 mg/ml L-rhamnose respectively, cultured for 8 hours and fluorescence measured by flow cytometry. Error bars shown in both A and B represent the standard deviation of the mean of three independent biological replicates.

As reporter-GFP mRNA molecules from the multicopy plasmid will not be produced in any gene silencing application of sRNAs in Azospirillum, we suggest that full suppression of the reporter mRNA at this screening stage is unnecessary. However, to confirm that sRNAs were capable of full mRNA silencing, another sRNA-reporter pair targeting *tetR* and previously used in *E. coli* ^41^, was introduced into pER100, resulting in plasmid pER158. Subsequently the L-rhamnose inducible promoter was replaced with a strong constitutive promoter identified through the SPL experiments (pER419), resulting in plasmid pER303. Both plasmids were tested in *A. brasilense*. As before, a partial reduction in reporter-GFP fluorescence was observed in cells containing the L-rhamnose promoter version. But consistent with our hypothesis that increased sRNA transcription would lead to full translation inhibition, no fluorescence was observed in cells with the constitutive promoter version (Figure 5B), indicating that pER100 is a useful screening plasmid for sRNA function and that full inhibition of reporter-GFP translation is not required to take forward for future sRNA applications.

### Engineering *A. brasilense* strains for the overproduction of the important plant-growth promoting hormone, indole 3-acetic acid (IAA)

As an initial demonstration of the power of the suite of genetic tools for transcriptional and post-transcriptional regulation developed for Azospirillum, we aimed to produce overproduction strains of IAA, a key plant stimulant. These strains could be used to purify the compound which is subsequently added to seeds or crops in the field; or they could be used directly as microbial biostimulants for plant-growth promotion in commercial agriculture. To do this we wanted to test promoters with a range of expression strengths, from weak to very strong, in combination with the native *ipdC* gene from A. brasilense, encoding the key enzyme in the production of IAA. We chose the Start-Stop Assembly^42^ one-pot assembly system for this, as it is straightforward to make our promoters, terminators and expression plasmid compatible with the system. Level 0 plasmids containing the promoters from pER407, 411, 416 and 419 were constructed. A level 0 plasmid containing a composite part with the *ipdC* gene and its native RBS from the *A. brasilense* genome were also constructed. The *E. coli* terminator, ECK120015170, validated in *A. brasilense* with pER206, was already contained in the original level 0 Start-Stop plasmid pGT417, so this was used in this work. Finally a new level 1 assembly plasmid, pER220, was constructed, combining the pER100 backbone containing the stable pBHR1 origin, and the appropriate SapI sites and *lacZ* expression cassette from pGT401. Level 1 *ipdC*-expression plasmids were constructed with each promoter (from pER406, 407, 411, 416, 419) and sequence verified by Sanger sequencing. *A. brasilense* cells were transformed with the resulting plasmids, pER221-225 and cultured in 50 ml cultures of LB supplemented with 3 g/L L-tryptophan. When cells reached stationary phase at an optical density of 1 (after approximately 18 h), they were normalised by dilution into fresh media and the supernatant isolated by filtration. The supernatant was subsequently mixed 1:2 with Salkowski’s reagent, incubated at room temperature for 30 mins and the concentration of IAA quantified by monitoring absorbance at 530 nm^43^. Using a standard curve of purified IAA in media, the mean concentration of IAA produced by each strain was calculated and plotted (Figure 6). *A. brasilense* naturally produces IAA when grown in rich media with tryptophan as seen with cultures containing the empty vector pER220. However when analysed using a one-way ANOVA (p-value: <0.0001) and t-tests with Tukey’s correction for multiple comparisons, all cultures containing the promoter-*ipdC* expression plasmids showed significantly-higher differences in mean IAA concentrations to the empty vector control cultures (p-values: 0.0013, <0.0001, 0.0001, <0.0001, <0.0001 respectively). The pattern of increasingly strong promoters that was observed in the SPL experiments, corresponded to increasing IAA concentrations with the exception of pER223-containing cultures. The strain containing the plasmid with the strongest promoter, pER225, produced almost three times as much IAA as the native wild-type strain.

**Figure 6.**
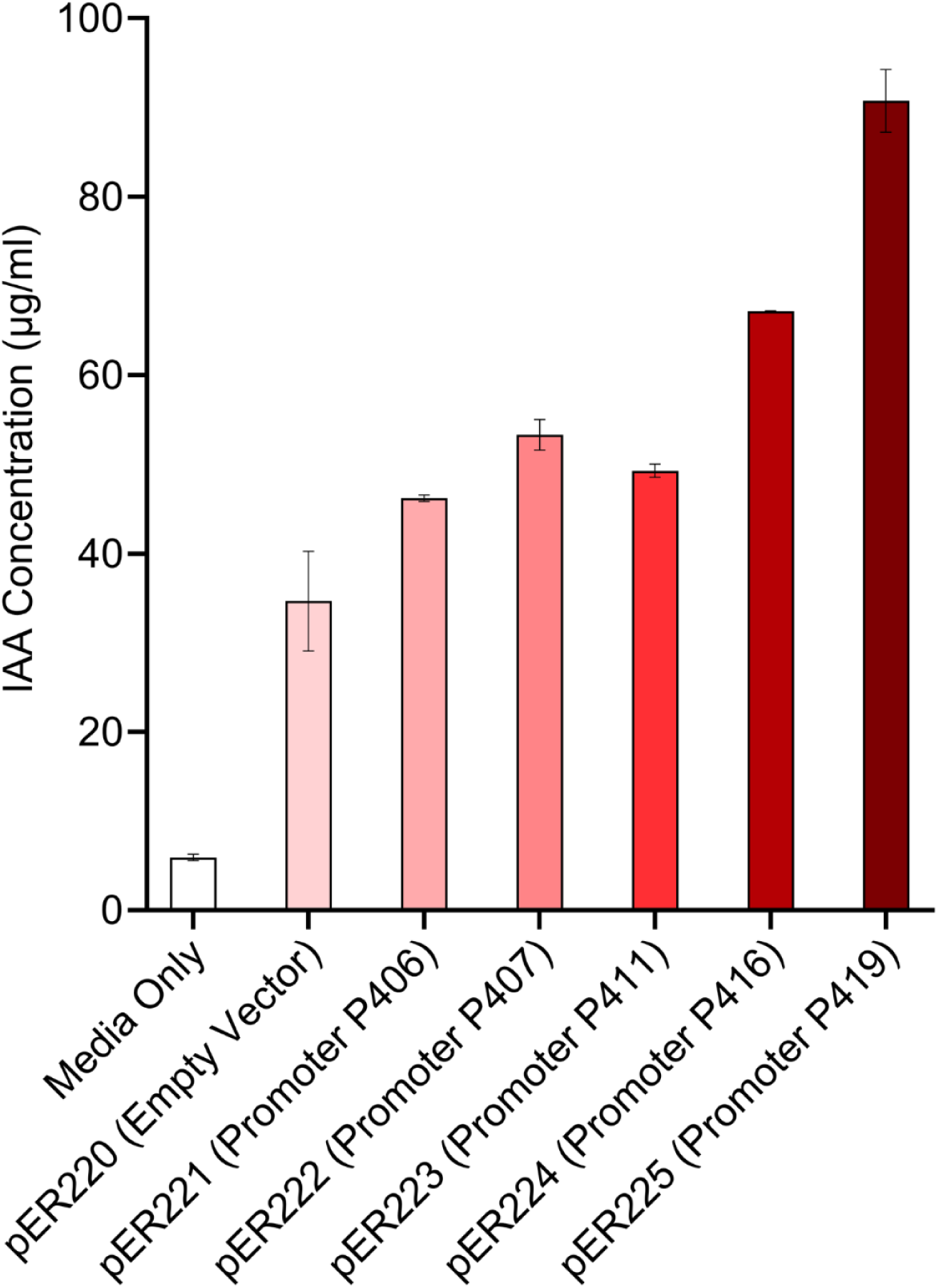
Overproduction of indole 3-acetic acid (IAA) in A. brasilense. Five /pdC-expression plasmids were constructed with promoters from plasmids pER406, 407, 411, 416, 419, resulting in plasmids pER221-225. *A. brasilense* cells were transformed with the resulting plasmids, and cultured in 50 ml cultures of LB media supplemented with 3 g/L L-tryptophan until they reached stationary phase (after approximately 18 h, at OD 1). Cells were normalised by OD through dilution with fresh media, the supernatant isolated by filtration and the concentration of IAA quantified using Salkowski’s reagent and monitoring absorbance at 530 nm. Error bars represent the standard deviation of the means of three independent biological replicates.

## Conclusion

In this work, we have developed the first comprehensive toolkit of regulatory genetic parts allowing high-resolution control of gene expression in any species of the important agricultural bacterial genus, *Azospirillum*. We have created a diverse set of synthetic constitutive promoters based on the consensus promoter sequences of *A. brasilense* sp7, with a wide range of expression strengths, including some with much higher expression levels than previously reported. We have adapted and validated a robust inducible promoter with low basal expression, a non-metabolisable inducer and one that does not require heterologous expression of the transcription factor, reducing the burden often associated with systems requiring the expression of a non-native transcription factor. We have introduced post-transcriptional regulation through the validation of synthetic sRNAs, which can be used for gene knockdown and/or gene silencing to be achieved, as well as the construction of multi-level (DNA-RNA-protein) circuits for more sophisticated programming of cells in an agricultural setting. We developed a one-pot sRNA/reporter cloning system to enable the rapid testing of sequences in *Azospirillum* and other organisms. We have adapted a set of our promoters and terminators for use in the convenient Start-Stop assembly method^42^, as well as a new compatible expression plasmid. Finally, with no optimisation in either the growth process or metabolic engineering, we were able to generate IAA-overproducing strains that accumulate IAA at concentrations three times higher than the wild-type strain. It is likely that modifications such as the silencing of competing native pathways, changes to media formulations and bioreactor optimisation, the yields of these IAA overproducers can be increased to commercially-viable levels.

This work introduces *A. brasilense* as a 21st Century Engineering Biology chassis, bringing new possibilities to the area of agricultural biologicals as a precision microbial additive to seeds and soils. These tools could be used to: overproduce key hormones involved in controlling the growth versus defence trade-off that limits crop productivity in agricultural settings; introduce novel desirable compounds for crop growth and fight off phytopathogens; improve studies of plant-microbe interactions in the rhizosphere and indeed manipulate this microbiome; and finally, introduce the ability to programme a robust commercial agricultural bacterium with new complex biocomputation to allow it to sense and respond to environmental challenges facing crops.

## Materials and Methods

### Bacterial strains and general laboratory practice

All bacterial strains and plasmids used in this study are listed in the Supplementary Information Table S1 and S2. *A. brasilense* sp7 was obtained from DSMZ (DSM1690), aerobic growth was achieved by culturing bacteria in Luria Bertani (LB) medium or in minimal medium for *Azospirillum brasilense* (MMAB) at 30 °C with orbital shaking at 200 rpm. MMAB media: K2HPO4 (3 g/L), NaH2PO4 (1 g/L), NH4Cl (1 g/L), MgSO4 7H20 (0.3 g/L), KCl (0.15 g/L), CaCl2 2H20 (0.01 g/L), FeSO4 7H20 (0.0025 g/L), sodium malate (5 g/L) and biotin (0.005 g/L). Aerobic growth of *E. coli* strains was achieved by culturing cells in LB medium at 37 °C, with orbital shaking at 200 rpm. For growth on solid media, LB containing 1.5% agar was used and incubated at 37 °C static incubation. *E. coli* strain DH10β was used for all plasmid cloning, propagation and isolation in this study. Plasmids were extracted and purified using Thermo Scientific GeneJET Plasmid Extraction kit according to manufacturer’s instructions and sequence verified using Dundee DNA Sequencing and Services. Plasmid transformation with *E. coli* DH10B was performed by heat-shock according to NEB’s transformation protocol. Media was supplemented with antibiotics to the following final concentrations: carbenicillin (*A. brasilense*: 25 ug/ml, *E. coli*: 125 ug/m) and kanamycin (*A. brasilense and E. coli*: 50 ug/ml).

For experiments conducted in deep well plates, cells were cultured in MMAB overnight with appropriate antibiotic concentration. Subcultures were then normalised to an OD600 of 0.05 in fresh media and supplemented with the appropriate inducer concentration and grown at 30 °C with rapid shaking at 700 rpm until a desired cell-density was reached (for fluorescent assays this was late-exponential phase, OD600 0.8, and stationary phase). For microplate reader experiments, growth conditions were the same, but the final normalisation takes place in a shallow 96-well microplate which was then incubated in a Tecan microplate reader at 30 °C with rapid orbital shaking (800 rpm).

### Electroporation of *A. brasilense* sp7

For the preparation of electrocompetent cells, *A. brasilense* sp7 was revived from-80 °C glycerol stocks on LB agar containing 25 µg/ml carbenicillin and incubated for 36 hours at 30 °C. A single colony was picked to inoculate 5 ml of LB media containing 25 µg/ml of carbenicillin and incubated overnight1 ml of overnight culture was used to inoculate 100 ml of fresh LB media containing 25 µg/ml of carbenicillin and grown to an OD600 of 0.8. The culture was then transferred into two 50 ml sterile falcon tubes and left on ice for 30 minutes. The cultures were then pelleted by refrigerated centrifugation for 10 minutes at 4,000 x g, the supernatant was discarded, and the pellet resuspended gently in 10 ml of ice-cold double-distilled autoclaved water (ddH_2_0). A second centrifugation was performed, the supernatant was discarded, and the pellet was resuspended in 5 ml of ice-cold ddH_2_0. A final centrifugation step was carried out, the supernatant discarded, and the final pellet resuspended in 1 ml of ice-cold ddH_2_0. The final resuspension was aliquoted in volumes of 200 µl and stored at-80 °C. For transformation with plasmid DNAprepared cells were thawed on ice and 0.2 cm cuvettes were chilled in the freezer for approximately 30 minutes. Pre-prepared LB media and LB agar plates containing selective antibiotics were pre-warmed in a water bath or 30 °C incubator respectively. To 40 µl of thawed cells 500 ng/µl of purified plasmid DNA was added and gently mixed on ice. The mixture was then added to the pre-chilled 0.2 cm cuvettes and tapped three times to remove bubbles. Cuvette electrodes were then wiped with tissue paper to remove any excess liquid. Electroporation was performed at 2.5 kV using the defaults settings in a Bio-Rad MicroPulser electroporator (Bio-Rad Laboratories). Immediately after electroporation cells were resuspended in 950 µl of pre-warmed LB media, transferred into a 1.5 ml Eppendorf and recovered at 30 °C with shaking at 200 rpm for 2 hours. Following recovery, 50 µl of cells were plated on selective LB agar plates.

### Plasmid stability assays

To determine plasmid stability in *A. brasilense* sp7 cultures, a modified version of the colony patching method^44^ previously reported was used. Electrocompetent *A. brasilense* sp7 cells were transformed with individual plasmids pBHR1 (pBHR ori) and pCK405 (p15a ori) on relevant selection LB agar plate. Single colonies were isolated and grown in triplicate overnight in 5 ml of LB media with plasmid-relevant antibiotics. The overnight cultures were passaged by diluting 1:1000 into 5 ml of fresh LB media containing no antibiotic and grown for 24 hours at 30 °C with shaking at 200 rpm. After 24 hours of cultivation 1 ml of sample was taken from each culture, centrifuged at 12,000 rpm, the supernatant discarded, and the pellet resuspended in 100 µl of 1X phosphate buffered saline (PBS). The resuspension was then serially diluted to 1:10,000 in PBS and plates onto LB agar plates containing no antibiotics and selective antibiotics in parallel, plates were incubated at 30 °C overnight. These passaging steps were repeated for a total of five passages. Enumeration of plasmid loss was calculated by counting the colony forming units (CFU) on selection and non-selection plates to check for colonies with lost antibiotic resistance (indicating plasmid loss). The % of plasmid loss was calculated by dividing the CFU on antibiotic plates by the CFU on no antibiotic plates and multiplying by 100.

### Assembly of the synthetic promoter library

We used a simple bioinformatic pipeline (Poma et al. unpublished) to identify the consensus-35 and-10 promoter sequences from the reference genome of *Azospirillum brasilense* sp7 (NZ_CP012914.1) available on NCBI, BLAST. To identify native promoter sequences, the genome was divided into six sequences for ease of handling. BPROM software was used to obtain total predicted promoter sequences in *A. brasilense* sp7 (NZ_CP012914.1) genome. Characters were removed from the resulting nucleotide motifs in Sequence Massager, and an original Python script (Poma et al. unpublished) was used to isolate the-35 and-10 promoter sequences from the ORF-less genome (Figure 2A). A sequence logo of the total isolated-35 and-10 promoter sequences was generated with WebLogo software, where heights of nucleotide symbols within the stack indicate relative frequency of nucleic acid at that position (Figure 2B) ^11^. With the relative nucleotide frequencies for the-35 and-10 promoter sequences in *A. brasilense* sp7 obtained, we determined thresholds of nucleotide conservation with the aim to maintain true levels of conservation but prevent the escape of sequence diversity. The final consensus sequence was the template for mutagenic primer design and synthetic promoter library construction by introducing sequences upstream of a gene encoding *sf*GFP on the pBHR1 plasmid backbone, using degenerate oligonucleotides, Q5 PCR amplification and blunt-end ligation. After propagation and screening in *E. coli* cells, *A. brasilense* sp7 cells were transformed with this library; 94 single colonies were cultured in microplates until late-exponential phase of growth, and *sf*GFP fluorescence measured using a microplate reader to determine the preliminary range of expression strengths present in the library. Following confirmation that the library contained phenotypic variation, twenty clones were isolated and tested in triplicate in fluorescence assays using flow cytometry

### Assembly and testing of the L-rhamnose inducible promoter system

For the construction of the l-rhamnose inducible plasmids pER70 (*rhaS*) and pER72 (no *rhaS*) the P*rhaBAD* promoter was inserted upstream of the *A. brasilense* optimised RBS sequences by blunt-end ligation. The *rhaS* activating transcription factor was inserted downstream of the kanR gene in pER72 by gibson assembly. Prior to inducible promoter characterisation we conducted carbon metabolism assays in *A. brasilense* sp7 to identify potential non-metabolizable inducer candidates. *A. brasilense* sp7 cultures were grown in MMAB media without its preferred malate carbon source, instead cultures were grown in triplicate with 0.5% L-Rhamnose and 0.5% L-Arabinose respectively, in a time course plate reader experiment in appropriate culture conditions. *A. brasilense* sp7 transformants containing plasmids pER70 and pER72 cultured in MMAB media supplemented with kanamycin and increasing concentrations of L-Rhamnose (0.0-0.4 mg/ml). Plasmid pER35 represents the constitutive promoter control BBa_J23118. Fluorescence intensity was detected using flow cytometry at 8 h (after the culture was inoculated).

### Assembly and testing of terminator plasmids

Terminators were placed between the L-Rhamnose inducible promoter and the ribosome binding site of the *sf*GFP reporter gene on pER70 by Q5 site-directed mutagenesis and blunt-end ligation. *A. brasilense* sp7 transformants containing the appropriate terminator between the L-Rhamnose inducible promoter and the ribosome binding site of the sfGFP reporter gene plasmids alongside no promoter control (pER65) and no terminator control (pER70) were cultured in MMAB media supplemented with kanamycin and 0.4 mg/ml L-Rhamnose. Fluorescence intensity was detected using flow cytometry at 8 h (after the culture was inoculated).

### One-pot assembly method for sRNA and reporter sequences

A single broad-host range plasmid system that is stable in *A. brasilense* sp7 and allows the rapid assembly and screening of synthetic sRNAs against reporters was designed. A one-pot restriction-ligation assembly reaction was used for the construction of Level 0 and Level 1 plasmids used in this study. Level 0 reporter and sRNA sequences were introduced into the start-stop assembly vector pGT400 by PCR amplification and blunt-end ligation^42^. Level 0 reporter or sRNA sequences were designed with unique 5’ and 3’ SapI recognition sites for the downstream cleavage and ligation in the Level 1 entry vector pER100 at specific SapI restriction sites. Level 1 assembly reactions contained 20 fmol of the destination vector plasmid DNA, 40 fmol of each insert (Level 0 plasmid DNA), 2 µl of T4 DNA ligase buffer, 1 µl T4 DNA ligase and 1 µl SapI restriction endonuclease in a total reaction volume of 20 µl made up with nuclease free water. Reactions were incubated using a thermocycler for 35 two-step cycles of 37 °C for 5 minutes and 16 °C for 5 minutes, before a single final denaturation step at 65 for 20 minutes. The total reaction mixture was transformed with competent cells for further plasmid selection and verification.

### GFP-fluorescence assays

A microplate reader (Tecan) was used to screen the initial promoter library in *E. coli* DH10B strains and *A. brasilense* sp7. Fluorescence of GFP containing cells was measured using 485 nm and 535 nm as the respective excitation and emission wavelengths, and optical density measured by absorbance at 600 nm. Kinetic cycles were carried out for 15 – 24 hours. Raw data was obtained from normalised fluorescence calculated by dividing fluorescence by absorbance at each timepoint. Data analysis and visualisation was performed in Prism GraphPad using stationary phase data. Flow cytometry was used to detect the fluorescence of individual cells from populations containing the SPL, L-rhamnose inducible promoter system, terminator and synthetic sRNA and reporter plasmids used in this study. Flow cytometry was carried out using the FACSCanto™ II High Throughput Sampler (HTS) and the BDS software to observe cell distribution and fluorescence in cell populations transformed with plasmids used in this study. Fluorescence intensity of sfGFP was detected using a 450 V blue laser, size and granularity of cells was measured using the Forward Scatter (FSC) laser at 413 V and the Side Scatter (SSC) laser at 334 V. The injected sample volume was 50 µl and the sample flow rate was 1.5 µl/sec, 10,000 events were recorded per sample. Raw data exported from the BDS-FACSCanto™ software was analysed using FCS Express flow cytometry software to obtain the geometric means of the populations under investigation. Data retrieved from FCS Express was analysed and visualised using GraphPad Prism software.

### sRNA-dependent translation inhibition assays

Electrocompetent *A. brasilense* sp7 cells were transformed with appropriate sRNA-reporter plasmid and plated onto LB agar containing 50 µg/ml kanamycin. Single colonies were isolated in triplicate and used to inoculate 5 mL of MMAB media and cultured overnight. Overnights were pre-cultured in 96-well deep plates containing 300 µl of fresh MMAB with 50 µg/ml kanamycin before incubation at 30 °C with rapid shaking at 700 rpm for 5 hours. Pre-cultures were then normalised to an OD600 of 0.05 by dilution into fresh media and supplemented with the appropriate l-rhamnose concentration and grown at 30 °C with rapid shaking at 700 rpm until a desired optical density was reached (this was late-exponential phase, OD_600_ 0.8, and stationary phase). Cultures were then diluted 1:1000 into sterile PBS before fluorescent quantification by flow cytometry.

### Design of *A. brasilense* sp7 synthetic sRNAs and reporter sequences

To design synthetic sRNAs targeting *mutL* and *mutS* in *A. brasilense sp7*, the reference genome for *A. brasilense sp7* (NZ_CP012914.1) was downloaded from NCBI. The reverse complement of various 24 bp seed regions covering the 5’UTR and the Shine-Dalgarno sequence of the target mRNA sequence were designed using SnapGene’s primer design tool and analysed using the Freiburg RNA Tool: IntaRNA ^40^. Query sRNA sequences were analysed for binding to the ribonucleome of the *A. brasilense sp7* (NZ_CP012914) reference genome using the Freiburg RNA Tool: IntaRNA default settings. IntaRNA’s algorithm uses RNA secondary structure predictions to predict the combination of the free energy required for accessibility to the RNA-RNA interaction site – in this case assessing the affinity of synthetic sRNAs for target mRNA sequences.

### *ipdC*-overexpression plasmid assembly

Plasmid pER220 was built using Gibson Assembly^45^ combining the backbone from the broad-host range pER100 and the start-stop compatible *lacZ* insert from pGT401 and appropriate SapI restriction sites. Plasmids pER221-225 were constructed by Start-Stop assembly of this new pER220 level one plasmid, and level 0 plasmids containing promoters, the RBS-*ipdC* gene and terminator.

### Indole-3-acetic acid quantification

*A. brasilense* sp7 cells were transformed with plasmids, pER220-225, and cultured in the dark in 50 ml cultures of LB supplemented with 3 g/L L-tryptophan and kanamycin. When cells reached stationary phase at an optical density of 1 (after approximately 18 h), they were normalised by dilution into fresh media and the supernatant isolated by filtration. The supernatant was subsequently mixed 1:2 with Salkowski’s reagent, incubated at room temperature for 30 mins in the dark and the concentration of IAA quantified by monitoring absorbance at 530 nm ^43,46^ Using a standard curve of purified IAA in media, the mean concentration of IAA produced by each strain was calculated and plotted. IAA and L-Tryp stock solutions were prepared according to previously-reported studies ^47^.

## Acknowledgments

This work was supported through a ESCMID Research Grant 2020 (CK) and the NERC OnePlanet DTP (ER).

