## Supplementary Tables and Figures for "Advancing genetic engineering in the plant growth-promoting agricultural bacterium *Azospirillum brasilense*"

### Supplementary Tables for Materials and Methods

**Table S1 of synthetic promoter sequences used in this study.**

| Plasmid | Promoter sequence 5' 3' |
|---|---|
| pER400 | TTTCCAGGAGGCCCCACCGGGTTCCCTCTCCAAACAAATTTGTTT |
| pER401 | GGTCTGACGCTTGAATACGTTGAAGTAGTCCGTTAACATAGCCAT |
| pER402 | ATACATGTCAACGATAATACAAAATATAATACAAACTATAAGATG |
| pER403 | TCGCACTTGCTTGACATACGGGAACTAGGCTGAGATA |
| pER404 | AAATAACCATTTGATTCCTACGCATGGCCACACTCCTCAT |
| pER405 | CAGAGAAATTTTATGTCACAGTACGTTGTGGCCACTATCGACTC |
| pER406 | AAAGCATATTGACTACGAGCGCTAGCTGTAACATGATGCGCCTTA |
| pER407 | AATGGAATGATTGCATTTTAAAACTAATATCCGTAAACTTGCAAA |
| pER408 | GGGAGACCACAACGGTTTCCCTCTACAAATAATTTTGTTTAACTT |
| pER409 | TCGATGCCAGTTGACAATCTTTCCTAGCTTAATTATGCTTACATA |
| pER411 | CCCAATTATTTGAAGTATTTTTCGGTACTTACTTTATTAT |
| pER412 | CGAGTAAGCATTGCTAAGATAGGCGATGCTGTATAGAATCTATAA |
| pER413 | AACCGCCACGTTGACTGGGAAGGGTGACTTGGCATATCATATCGAT |
| pER414 | CCACAATTCAGCACATTGTGAACATCATCACGTTCATCTTTCCCT |
| pER415 | CTTCACACCTTGACAACGAAGTCTATAGTTTACATACTTTTAGTT |
| pER416 | TGTTCACCCCAACGCATTCCCTTTTCTAGTAAACATACTTTTAGT |
| pER417 | GCGCATTTGATTGAAGAGAGTATCCATGTGGACTAAGCTAGGCAG |
| pER418 | GGCCTGCGATTTGACAACGAGGGCAATGGGAGTCATTCTGAGATT |
| pER419 | TGCGTTTATATTGACGGATATCTCAGTCCTAGGTATTGTGCTCGT |

| | |
|---|---|
| pER420 | TCTTTGCTTGCTTTTTTACACTTTATGCTTCCGGCTCGTATGTTG |

**Table S2 of terminator sequences used in this study**

| Plasmid | Terminator Sequence 5' 3' |
|---|---|
| pER200 | GGAAACACAGAAAAAAGCCCGCACCTGACAGTGCGGGCTTTTTTTTTCGACCAAAGG |
| pER201 | CTCGGTACCAAATTCCAGAAAAGAGGCCTCCCGAAAGGGGGGCCTTTTTTCGTTTTGGTCC |
| pER202 | GGAAACACAGAAAAAAGCCCGCACCTGACAGTGCGGGCTTTTTTTTTCGACCAAAGG |
| pER203 | CTCGGTACCAAAAAAAAAAAAAAAAGACGCTGAAAAGCGTCTTTTTTCGTTTTGGTCC |
| pER204 | TTTTCGAAAAAAGGCCTCCCAAATCGGGGGGCCTTTTTTATTGATAACAAAA |
| pER205 | TACTGATTTTTAAGGCGACTGATGAGTCGCCTTTTTTTTGTCT |
| pER206 | ACAATTTTCGAAAAAACCCGCTTCGGCGGGTTTTTTTATAGCTAAAA |
| pER207 | AACGCATGAGAAAGCCCCCGGAAGATCACCTTCCGGGGGCTTTTTTATTGCGC |
| pER208 | |

**Table S3 of synthetic sRNA sequences**

| Query description | Query Sequence sRNA 3' 5' | Overall Energy Score | Target ID |
|---|---|---|---|
| *A. brasilense* sp7 Anti-mutS | UCAUGGGGUUAUGUCCGGUUCCCU | -40.36 kcal/mol | MutS |
| *A. brasilense* sp7 Anti-mutL | CGGCAUGCGCCAAAUAUAGUAGGC | -36.91 kcal/mol | MutL |

**Table S4 of reporter sequences**

| Reporter | Reporter sequence 5’ 3’ |
|---|---|
| MutS | GCCCGCCACAACGCAACCGGTTGAAGACCTGTGTCCGACGCGTCCGCCGCCACTTCGCCC |
| MutL | TTCCCGGCGTGAGCCTACTATATTTGGCGCATGCCGATCCGTATGCTGCCCGACACGCTC |
| TetR | TCGATAAACACAGAGAAGTAGGTTCAGAATGTCCAGATTAGATAAAAGTAAAGTGATT |

**Table S5 of Plasmids**

| Plasmid | Description | Antibiotic Resistance | Source |
|---|---|---|---|
| pBBR1MCS | Broad-host range vector with pBBR1 ori, rep and mob and multiple cloning sites | Cml | Palmer Lab |
| pLD1 | Lac operon cassette with BsaI sites, pBBR1 ori and rep | Kan | TP Lab |
| pER25 | pLD1 + removal of lac operon cassette via BsaI assembly and insertion of Level 1 assembly sRNA/reporter cassette containing: sacB, PrhaBAD, micC, T1/TE, J23118 promoter, lacZ and noRBS_sfGFP, term, with unique Sap1 sites (Figure 3.X) | Kan | This study |
| pER27 | pER25 + blunt-end ligation to remove the Level 1 assembly sRNA cassette leaving the J23118, no RBS, sfGFP sequence | Kan | This study |

| | | | |
|---|---|---|---|
| pER35 | pER27 + J23118 + blunt-end ligation to insert Azo synthetic RBS 1 (high TLR) between J23118 and sfGFP | Kan | This study |
| pER36 | pER27 + blunt-end ligation to remove J23118 and gibson assembly to replace with ProD promoter, Azo synthetic RBS 1 (high TLR) | Kan | This study |
| pER65 | pER35 + blunt-end ligation to remove J23118 constitutive promoter, used as no promoter control in terminator characterisation experiments | Kan | This study |
| pER72 | pER35 + blunt-end ligation to remove J23118 promoter and replace with PrhaBAD promoter sequence | Kan | This study |
| pER70 | pER70 + gibson assembly to insert synthetic RBS + rhaS g block downstream of KanR | Kan | This study |
| pER200 | Broad-host range plasmid containing pBHR1 ori, J23118 promoter, Terminator ECK120033737; thrL attenuator, *A. brasilense sp7* optimised RBS 1 and sfGFP | Kan | This study |
| pER201 | Broad-host range plasmid containing pBHR1 ori, J23118 promoter, Terminator L3S2P21, RBS 1 and sfGFP | Kan | This study |
| pER202 | Broad-host range plasmid containing pBHR1 ori, J23118 promoter, Terminator X, RBS 1 and sfGFP | Kan | This study |

| | | | |
|---|---|---|---|
| pER203 | Broad-host range plasmid containing pBHR1 ori, J23118 promoter, Terminator X, RBS1 1and sfGFP | Kan | This study |
| pER204 | Broad-host range plasmid containing pBHR1 ori, J23118 promoter, Terminator X, RBS 1 and sfGFP | Kan | This study |
| pER205 | Broad-host range plasmid containing pBHR1 ori, J23118 promoter, Terminator X, RBS 1 and sfGFP | Kan | This study |
| pER206 | Broad-host range plasmid containing pBHR1 ori, J23118 promoter, Terminator X, RBS 1 and sfGFP | Kan | This study |
| pER207 | Broad-host range plasmid containing pBHR1 ori, J23118 promoter, Terminator X, RBS 1 and sfGFP | Kan | This study |
| pERSPL | pER35 template used for the one-pot blunt end ligation of the synthetic promoter library | Kan | This study |
| pER400 | Broad-host range plasmid containing pBHR1 ori, synthetic promoter, RBS 1, sfGFP | Kan | This study |
| pER402 | Broad-host range plasmid containing pBHR1 ori, synthetic promoter, RBS, sfGFP | Kan | This study |
| pER403 | Broad-host range plasmid containing pBHR1 ori, synthetic promoter, RBS, sfGFP | Kan | This study |

| | | | |
|---|---|---|---|
| pER404 | Broad-host range plasmid containing pBHR1 ori, synthetic promoter, RBS, sfGFP | Kan | This study |
| pER405 | Broad-host range plasmid containing pBHR1 ori, synthetic promoter, RBS, sfGFP | Kan | This study |
| pER406 | Broad-host range plasmid containing pBHR1 ori, synthetic promoter, RBS, sfGFP | Kan | This study |
| pER407 | Broad-host range plasmid containing pBHR1 ori, synthetic promoter, RBS, sfGFP | Kan | This study |
| pER408 | Broad-host range plasmid containing pBHR1 ori, synthetic promoter, RBS, sfGFP | Kan | This study |
| pER409 | Broad-host range plasmid containing pBHR1 ori, synthetic promoter, RBS, sfGFP | Kan | This study |
| pER411 | Broad-host range plasmid containing pBHR1 ori, synthetic promoter, RBS, sfGFP | Kan | This study |
| pER412 | Broad-host range plasmid containing pBHR1 ori, synthetic promoter, RBS, sfGFP | Kan | This study |
| pER413 | Broad-host range plasmid containing pBHR1 ori, synthetic promoter, RBS, sfGFP | Kan | This study |
| pER414 | Broad-host range plasmid containing pBHR1 ori, synthetic promoter, RBS, sfGFP | Kan | This study |

| | | | |
|---|---|---|---|
| pER415 | Broad-host range plasmid containing pBHR1 ori, synthetic promoter, RBS, sfGFP | Kan | This study |
| pER416 | Broad-host range plasmid containing pBHR1 ori, synthetic promoter, RBS, sfGFP | Kan | This study |
| pER417 | Broad-host range plasmid containing pBHR1 ori, synthetic promoter, RBS, sfGFP | Kan | This study |
| pER418 | Broad-host range plasmid containing pBHR1 ori, synthetic promoter, RBS, sfGFP | Kan | This study |
| pER419 | Broad-host range plasmid containing pBHR1 ori, synthetic promoter, RBS, sfGFP | Kan | This study |
| pER420 | Broad-host range plasmid containing pBHR1 ori, synthetic promoter, RBS, sfGFP | Kan | This study |
| pCK405 | Backbone for m-toluic acid inducible sRNAs, Pm promoter, tetR sRN, micC, T1/TE, p15a ori | Cml | Kelly lab |
| pCK302 | Backbone for L-Rhamnose inducible reporters, PrhaBAD, synthetic RBS, rrnB1 B2 T1 terminator, pBR322 ori | Amp | (Kelly et al. 2016) |
| pER1 | pCK405 + blunt- end ligation to insert the *E. coli* anti-*mutH* sRNA | Cml | This study |
| pER2 | pCK405 + blunt- end ligation to insert the *E. coli* anti-*mutS* sRNA | Cml | This study |

| | | | |
|---|---|---|---|
| pER3 | pCK405 + blunt- end ligation to insert the *E. coli* anti-*mutL* sRNA | Cml | This study |
| pER6 | pCK302 + blunt- end ligation to insert the *E. coli mutH*-sfGFP reporter | Amp | This study |
| pER7 | pCK302 + blunt- end ligation to insert the *E. coli mutL*-sfGFP reporter | Amp | This study |
| pER8 | pCK302 + blunt- end ligation to insert the *E. coli mutS*-sfGFP reporter | Amp | This study |
| pBBR1MCS | Broad-host range vector with pBBR1 ori, rep and mob and multiple cloning sites | Cml | Palmer Lab |
| pLD1 | Lac operon cassette with BsaI sites, pBBR1 ori and rep | Kan | TP Lab |
| pER25 | pLD1 + removal of lac operon cassette via BsaI assembly and insertion of Level 1 assembly sRNA/reporter empty module containing: sacB, PrhaBAD, micC, T1/TE, J23118 promoter, lacZ and noRBS_sfGFP, term, with unique Sap1 sites (Figure 3.X) | Kan | This study |
| pER100 | Level 1 empty vector, pER25 + Gibson assembly of mobilisation protein from pBBR1 MCS | Kan | This study |
| pER101 | Level 1 empty vector, pER100 + blunt-end ligation to remove terminator X and insert terminator Y | Kan | This study |

| | | | |
|---|---|---|---|
| pGT400 | Level 0 lacZ empty vector used for BsaI assembly of Level 0 synthetic sRNA and reporter plasmids. | Amp | (Taylor et al. 2009) |
| pER165 | pER100 + Sap1 assembly of the PrhaBAD anti-mutL/MutLsfGFP (*A. brasilense sp7* specific) synthetic sRNA screening combination | Kan | This study |
| pER158 | pER100 + Sap1 assembly of the PrhaBAD antitetR/tetRsfGFP synthetic sRNA screening combination | Kan | This study |
| pER303 | pER150 + blunt-end ligation to remove inducible PrhaBAD promoter and replace with constitutive synthetic promoter with the synthetic promoter sequence from pER419 (Table of plasmids 2.2. and table of promoter sequences 3.X) | Kan | This study |

**Table S6. Expression bin groupings of promoters.**

| Group | Log10_Fluorescence | Expression_Bin |
|---|---|---|
| pER35 | 3.73128 | Mid-Low |
| pER36 | 4.11547 | High |
| pER400 | 1.9893206666666667 | Low |
| pER401 | 2.1390403333333334 | Low |
| pER402 | 2.3130856666666664 | Low |
| pER403 | 2.322083333333333 | Low |
| pER404 | 2.3626056666666666 | Low |
| pER405 | 2.560069 | Low |
| pER406 | 2.6405246666666664 | Mid-Low |
| pER407 | 3.169909333333333 | Mid-Low |

| | | |
|---|---|---|
| pER408 | 3.690523666666667 | Mid-Low |
| pER409 | 3.7803210000000003 | Mid-Low |
| pER410 | 3.837358 | Mid-Low |
| pER411 | 3.8521983333333334 | Mid-High |
| pER412 | 3.897710333333334 | Mid-High |
| pER413 | 3.9411166666666664 | Mid-High |
| pER414 | 3.8842959999999995 | Mid-High |
| pER415 | 4.016143666666666 | Mid-High |
| pER416 | 4.248155333333333 | High |
| pER417 | 4.308606999999999 | High |
| pER418 | 4.373315333333333 | High |
| pER419 | 4.765976333333334 | High |
| pER420 | 4.7788216666666665 | High |
| pER65 | 1.7076736666666665 | Low |
| pER72 | 3.9680913333333336 | Mid-High |

#### Supplementary Figures

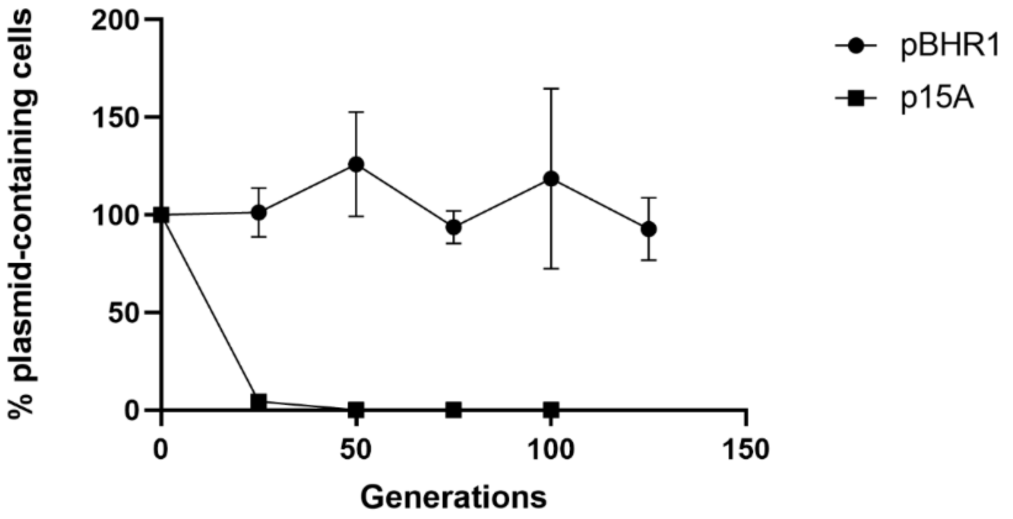


***Figure S1. Segregational stability of plasmids with two origins of replication in A. brasilense.***

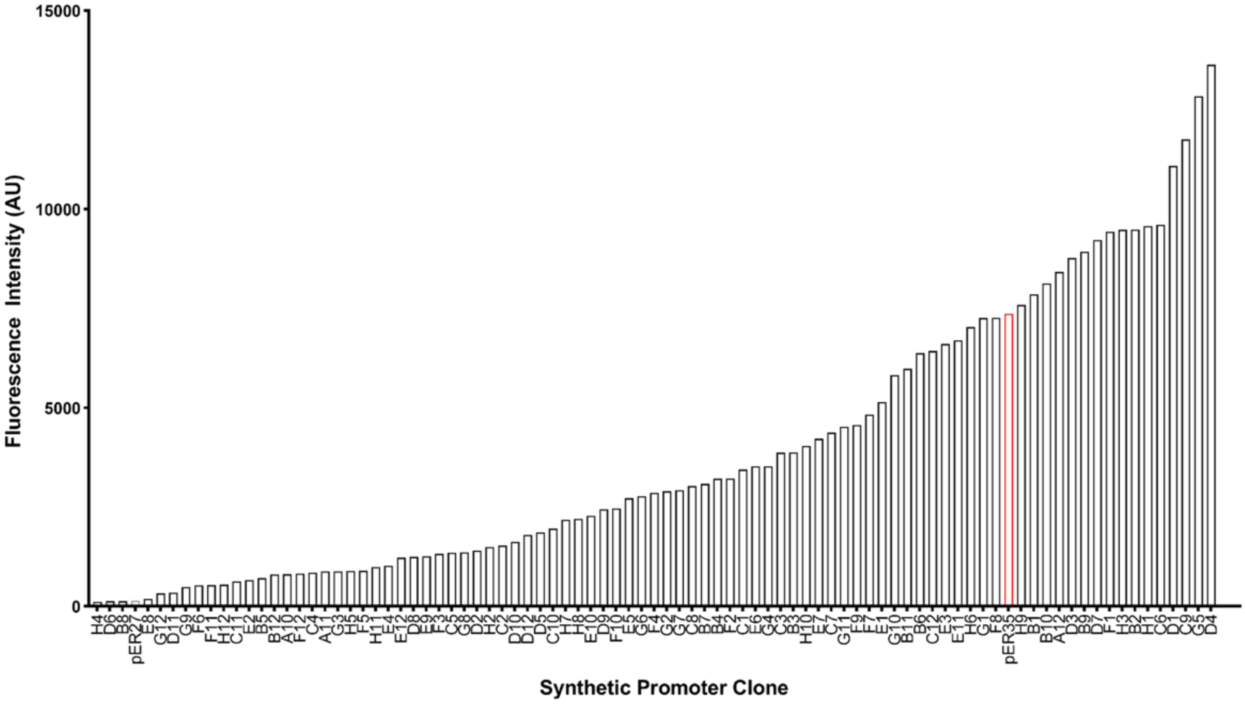


***Figure S2. Initial screen of SPL clones in A. brasilense.***

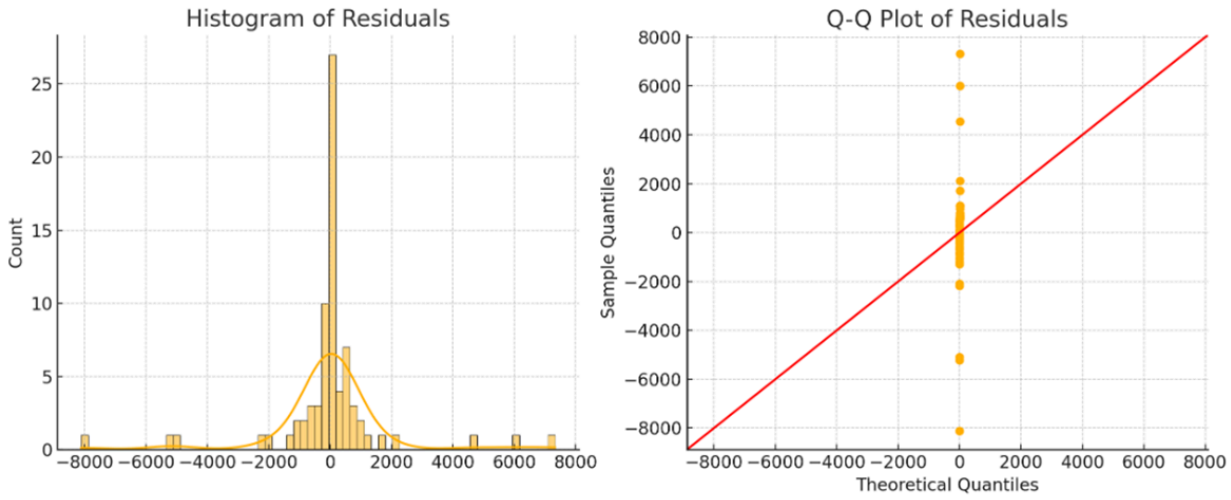


***Figure S3. Analysis of residuals from a fitted one-way ANOVA model of raw SPL mean fluorescence values.***

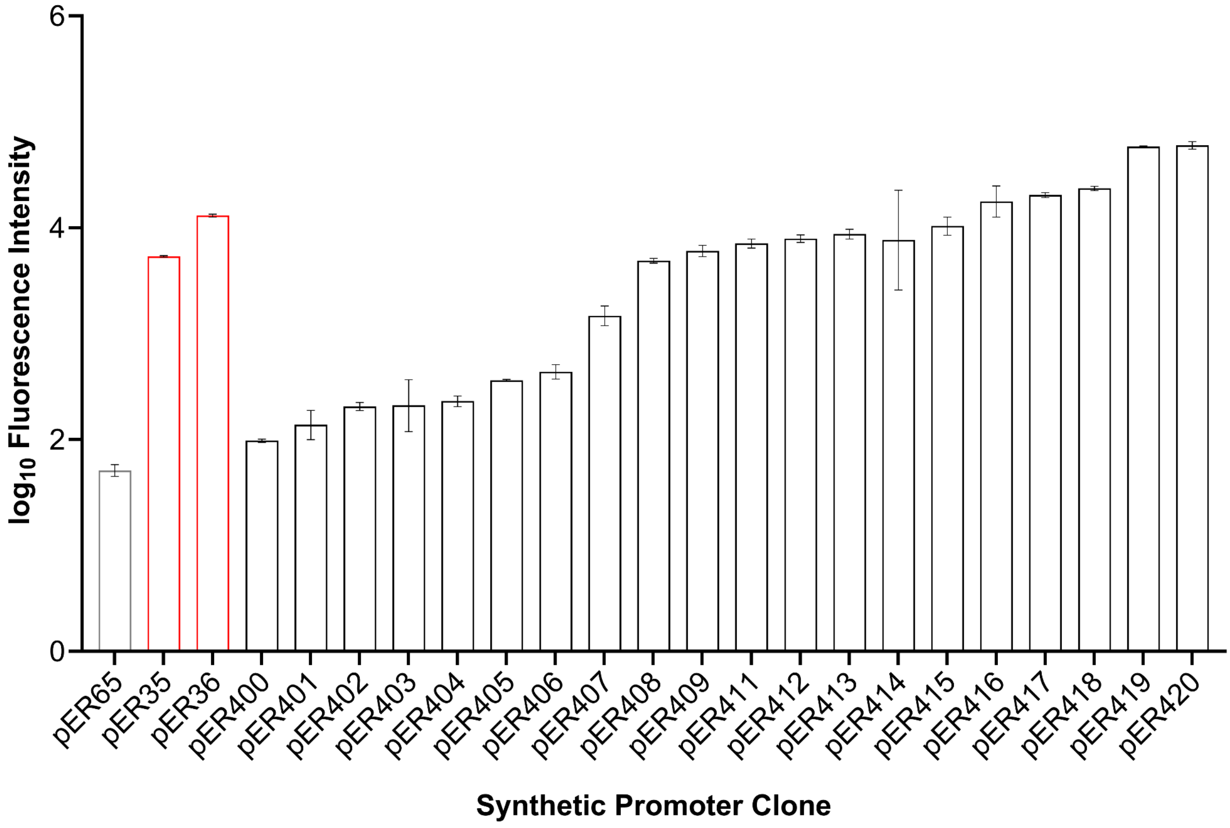


***Figure S4. log10 fluorescence values of SPL candidates.***

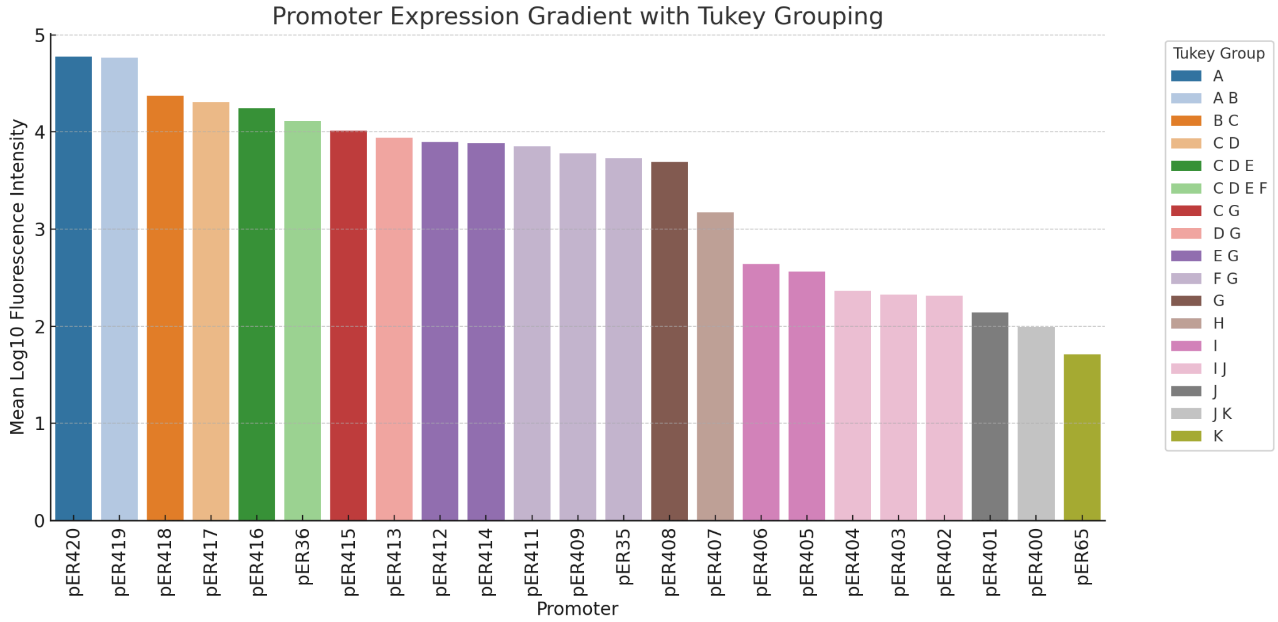


***Figure S5. Tukey grouping of promoter expression based on individual t-tests between each promoter, followed by Tukey's correction for multiple comparisons.*** A one-way ANOVA statistical test was conducted, showing a statistically-significant difference in the means across the promoter set (P = <0.0001). Individual t-tests with Tukey's corrections for multiple comparisons were conducted to assess pairwise differences between promoter means. All synthetic promoters except pER400 showed significantly-different fluorescence to the negative control plasmid pER65.

The results were summarized using a compact letter display, where promoters sharing a letter are not significantly different ($\alpha = 0.05$), while groups without shared letters are distinct.

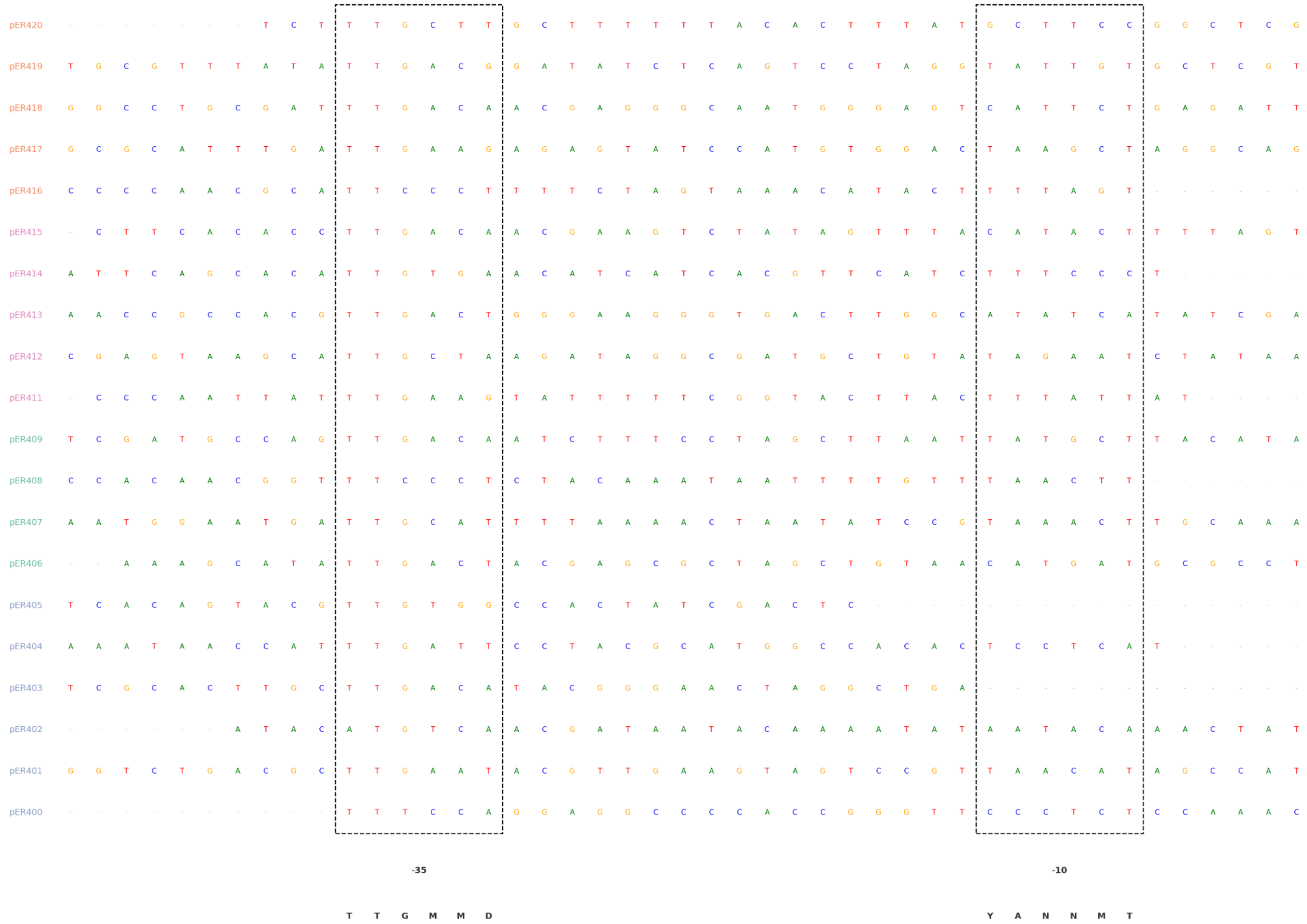


***Figure S6. Motif-anchored alignment of promoter sequences to consensus -35 and -10 operator sequences.*** Colour grouping of plasmid names represent expression strength bins: low to very strong from bottom to top.

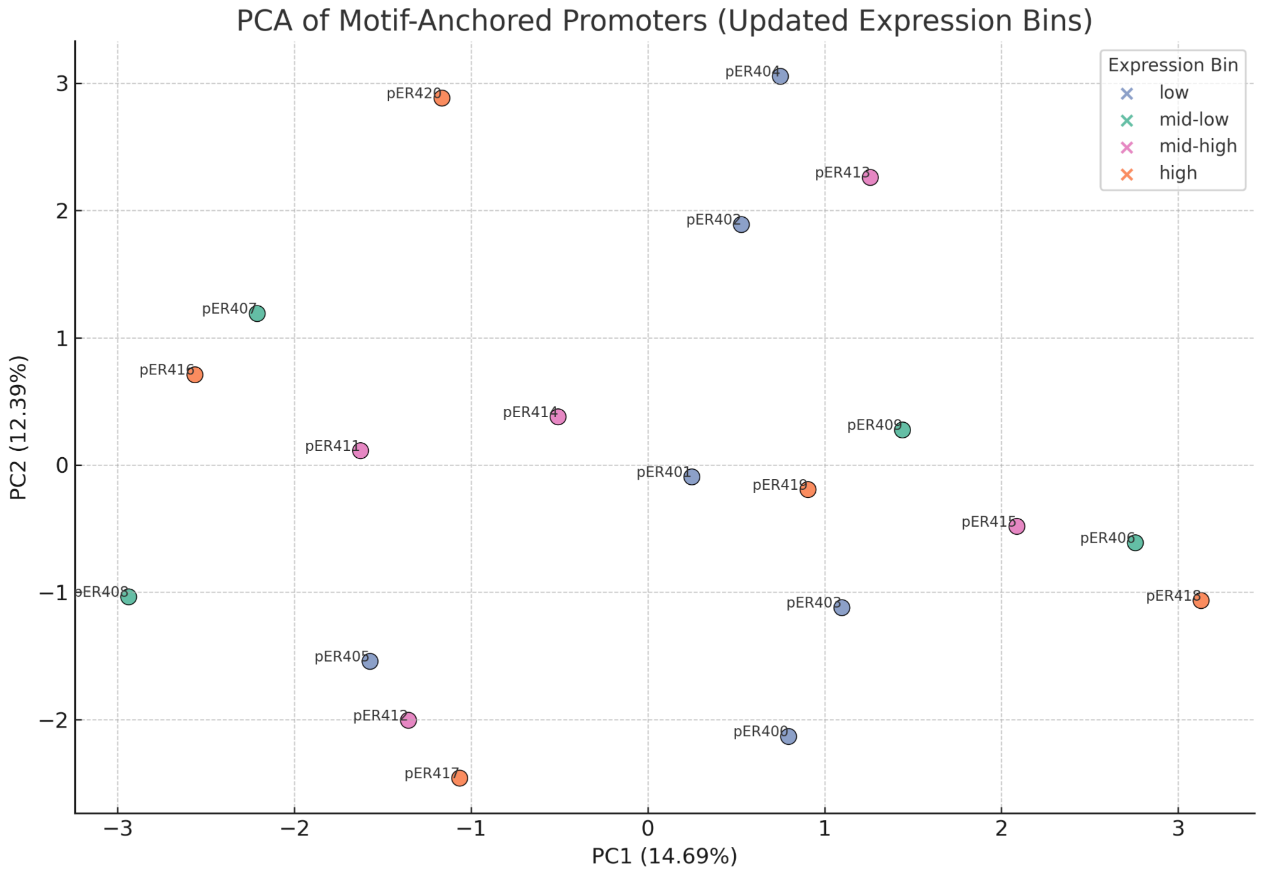


**_Figure S7. PCA of motif-anchored promoter sequence alignments._**

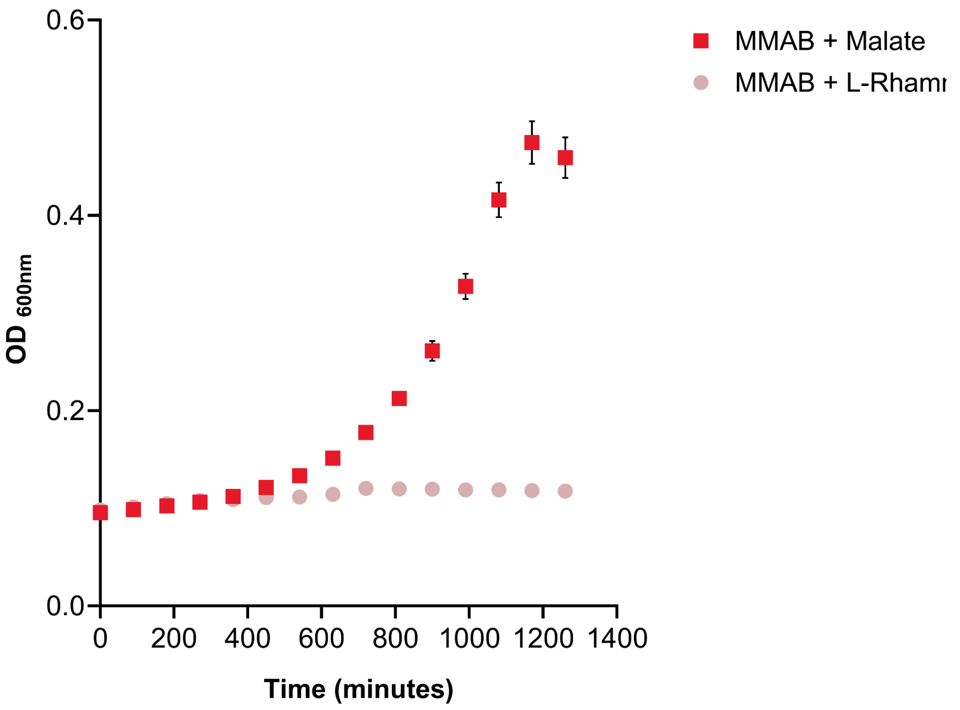


**_Figure S8. Growth curve analysis of A. brasilense cells cultured with malate or rhamnose._**

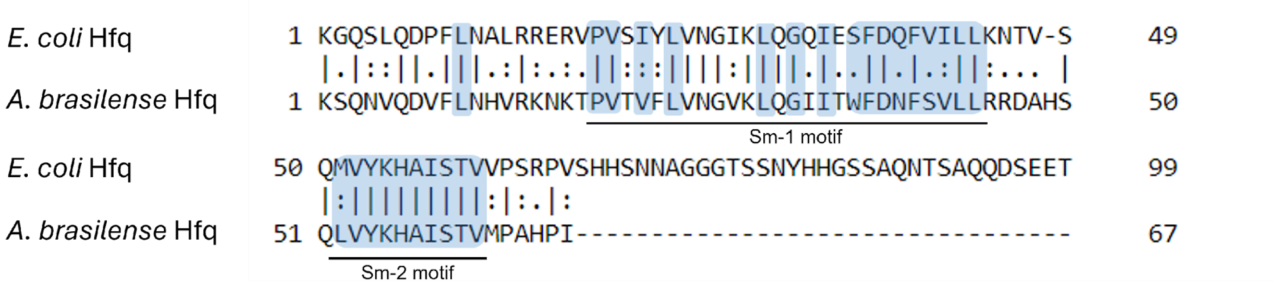


***Figure S9. BLAST alignment of E. coli and A. brasilense Hfq proteins.***
